# SyMBac: Synthetic Micrographs for Accurate Segmentation of Bacterial Cells using Deep Neural Networks

**DOI:** 10.1101/2021.07.21.453284

**Authors:** Georgeos Hardo, Maximilian Noka, Somenath Bakshi

**Affiliations:** Department of Engineering, University of Cambridge, CB2 1PZ, UK

## Abstract

We present a novel method of bacterial image segmentation using machine learning models trained with Synthetic Micrographs of Bacteria (SyMBac). SyMBac is a tool that allows for rapid, automatic creation of arbitrary amounts of training data, combining detailed models of cell growth, physical interactions, and microscope optics to create synthetic images which closely resemble real micrographs. The major advantages of our approach are: 1) synthetic training data can be generated virtually instantly, and on demand; 2) these synthetic images are accompanied by perfect ground truth positions of cells, meaning no data curation is required; 3) different biological conditions, imaging platforms, and imaging modalities can be rapidly simulated, meaning any change in one’s experimental setup no longer requires the laborious process of manually generating new training data for each change. Our benchmarking results demonstrate that models trained on SyMBac data generate more accurate and precise cell masks than those trained on human annotated data, because the model learns the true position of the cell irrespective of imaging artefacts. Machine-learning models trained with SyMBac data are capable of analysing data from various imaging platforms and are robust to drastic changes in cell size and morphology.

## Introduction

High-throughput time-resolved imaging of bacterial cells has revolutionised the fields of systems and synthetic microbiology [1, 2, 3, 4, 5, 6, 7]. Deep-learning based approaches are rapidly gaining popularity for automated processing of such high volumes of data [8, 9, 10, 11, 12, 13]. The performance of such deep-learning algorithms are fundamentally dependent on the quality and quantity of the training data provided to it [14]. However, the task of generating training data with accurate ground truth is not only slow and difficult but a near impossible one for images of micron-sized bacterial cells. The comparable size of the point spread function (PSF) of the microscope corrupts the images of bacterial cells, blurring them to the extent that neither expert users nor traditional computational segmentation programs can accurately annotate the correct pixels. Additionally, geometric impacts in the microscopic 2D projection of 3D objects in an image lead to inaccuracies in contrast based segmentation by both humans and computational programs (Supplementary Information 1). Inaccuracies in the ground-truth of training data cause deep learning models to misinterpret the relations between the object and its image, resulting in systematically artefactual mask predictions. This limits one’s ability to infer a cell’s true size and shape and confounds the analysis.

To address these limitations, we have developed SyMBac (Synthetic Micrographs of Bacteria), a tool to generate realistic synthetic images of bacteria growing in various imaging platforms. We combine understanding of cell growth and morphology, its position and interaction with the imaging platform, and the microscope’s optics to render synthetic images capable of training highly accurate segmentation convolutional neural networks (CNNs) without human annotated data. A neural network trained on this synthetic data can precisely learn about the corruption modes introduced during image formation, and as a result output accurate cell masks. This not only greatly speeds up the process of image segmentation (because no human annotation is necessary), but most importantly generates more accurate masks of cells, enabling precise analysis of size regulation and growth-dynamics. Using our method, given any change in experimental settings, it becomes trivial to generate high volumes of synthetic images and ground truths to rapidly analyse new data. This also addresses the robustness and reproducibility problems of image processing using deep-learning because models can now be easily adapted and benchmarked, since synthetic data can be generated at any desired spatial and temporal resolution with known parameters.

In this paper, we show that SyMBac can be used to generate synthetic micrographs for linear or monolayer colonies of bacteria in microfluidic devices or agar pads. While SyMBac can be easily used with fluorescence images, we primarily focus on the more challenging task of phase contrast images of bacteria in these platforms. This is because, phase contrast imaging can be used ubiquitously for all microbes since it does not require any kind of labelling, and also because it poses a more difficult challenge for segmentation algorithms, as the images contain the structures of the imaging platform making it difficult to separate them from the cells. This problem is most severely felt in linear colonies (mother machine [15]), and to a lesser extent single-cell chemostats [16], where the microfluidic device’s features are comparable to the size and refractive index of the cells which they trap. However, devices with linear colonies of bacteria are of major interest to quantitative microbiologists, as they are able to provide the highest throughput and most precise control of growth conditions [3], which have recently revolutionised research in quantitative single-cell bacterial physiology [15, 17, 18, 19, 20]. Accurate segmentation of individual cells in this platform is sorely needed for further and deeper investigations into singlecell physiology. Therefore, we focus on the linear colonies first, where the challenges of accurately identifying and segmenting individual cells is most sorely needed. Our results show that machine-learning models trained with SyMBac data produce more highly accurate masks of cells compared to models trained on data generated or curated by humans. This enables precise analysis of cell size and shape regulation along changing conditions in a growing bacterial cell culture, revealing novel insights about the physiological dynamics of individual bacterial cells during entry and exit from dormancy.

## Results

In Figure 1b, we show the overall process of generating synthetic training data for bacterial cells growing in linear colonies. A rigid body physics simulation is combined with an agent based model of bacterial cell growth (Supplementary Information 2). The output position and geometry of this combined model is then used to to rapidly produce many synthetic micrographs with ground truth. Properties from the simulation are extracted and used to render an optical path length (OPL) image that signifies the relative phase shift in the image—observed as as a change in intensity—due to the local refractive index (Supplementary Information 3, Figure SI 6). This image is then convolved with the microscope’s point spread function which can be generated using known parameters of the objective lens and phase ring/annulus (Supplementary Information 4). These properties are easy to find and thus any change in microscope optics can be readily simulated. The intensity distribution and noise spectrum of the convolved image is then optimised in order to maximise its similarity to the real image (Supplementary Information 5). Put together, this enables us to produce realistic phase contrast and fluorescence images of bacterial cells (Figure 1e and Figure 4). SyMBac takes advantage of multiprocessing, and where possible offloads array operations onto the GPU, which allows it to be approximately 10,000x faster than a human at generating training data (and if deployed on computing clusters, this could be extended by a further order of magnitude). Typical timescales of generating synthetic images are shown in (Figure 1d). The entire process of generating synthetic data has been made simple through the creation of example Jupyter notebooks with interactive widgets, which guide the user through the entire process.

**Figure 1:**
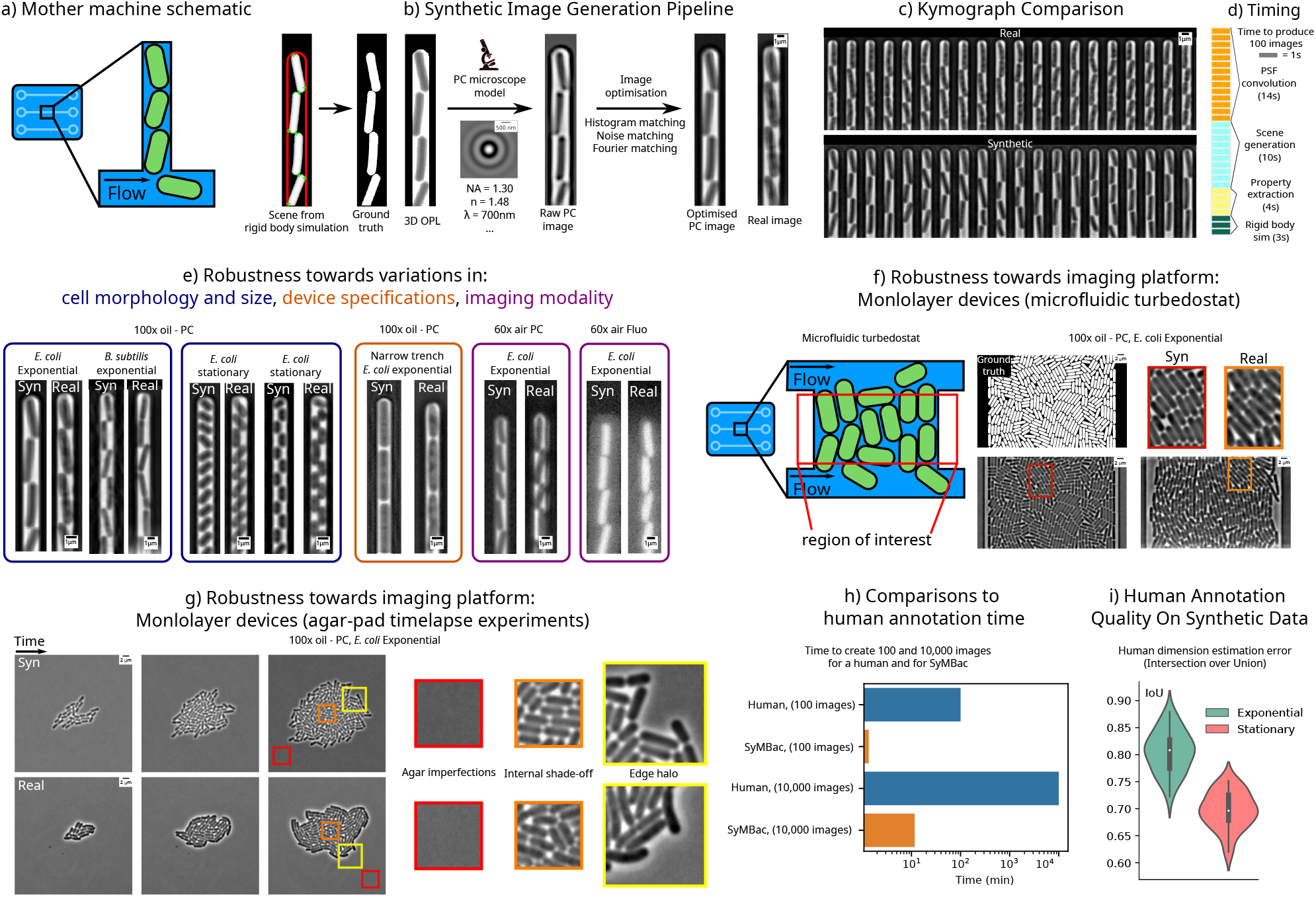
The synthetic image generation process: **a)** Schematic of linear colonies of cells in a microfluidic device known colloquially as the mother machine. **b)** Synthetic image generation pipeline: rigid body physics simulations are combined with agent based modelling to simulate bacterial growth in the device. These simulations are combined with the microscope’s point spread function, which is generated using known parameters of the objective lens. This output image is then further optimised to match real images. **c)** A kymogrpah of synthetic mother machine data generated by SyMBac, compared with a real kymograph from a timelapse experiment. **d)** Typical timescales for individual steps in the generation of training data. **e)** Synthetic data can be adapted to different biological conditions, variations in microfludic designs, and imaging modalities. With real data, many experiments would need to be conducted to generate training data with the same kind of coverage. **f)** Synthetic data can be generated for microfluidic devices that produce monolayer colonies, in this case the microfluidic turbedostat described in [16]. (Real image courtesy of Elf Lab, Uppsala University). **g)** SyMBac can also generate timelapse data for growth of monolayer colonies on agar pads. **h)** SyMBac is approximately 10,000x faster than a human at generating training data (10,000 images in less than 10 minutes) **i)** Humans annotating images had variable performances and consistently undersegmented cells, especially in small stationary phase cells.

Synthetic images generated by SyMBac can be tuned according to biology (cell size, shape, aspect ratio, optical path length), the imaging platform (microfluidic device geometry, agar pad), optics (objective lens), and whether the experiment is phase contrast or fluorescence (Figure 1e). The physics back-end of SyMBac can be adapted to model alternative imaging platforms, to generate synthetic training data for imaging experiments of both 1D and 2D colonies. In Figure 1f, we demonstrate the use of SyMBac to generate synthetic phase contrast images for the microfluidic turbidostat devices described in [16]. Comparison of the closeups from the synthetic image and the real image from such devices reveal the similarities in texture and contrast, which are crucial to ensure that the synthetic training data is realistic enough for models trained with it to perform well on real data (as shown in Figures 4, SI 20, SI 23, and SI 25). Similarly, it is possible to generate synthetic micrographs of timelapse experiments of the growth of monolayer colonies on agar pads. The close-ups of the synthetic images show that the model captures all the relevant details: background texture from agar imperfections, contrast shade-off effects near the centre of the monolayer colony, and background halo and darker cells near the edge of the colony (Figure 1g and Supplementary Information 11).

SyMBac data also enables benchmarking the performance speed and accuracy of human annotators and traditional thresholding based computational segmentation approaches (Figure 1). Our benchmarking experiments show that human annotators are not only slow, but generated inaccurate training data (Figure 1i), being unable to maintain consistency or accuracy in correctly identifying pixels at the cell boundary. Additionally we found that annotation results were highly variable across growth regimes, with the error rate in annotating small cells being exceptionally high (Figure 1). While better than human annotated training data, we also find that the unsupervised approaches using Otsu, local thresholding, and membrane dye images still suffer from bias and lack of robustness towards size variation (Figure SI 2, SI 3, and SI 4 in 1). This suggests that these traditional approaches are not suitable when highly accurate cell masks are desired, and would not even be suitable for generating training data required for ML algorithms to accurately segment data. SyMBac addresses this critical issue; since the synthetic images from SyMBac are accompanied with perfect ground truth information of cell shape and position. Models trained on our data therefore ensure the neural network learns the true relation between object and its corresponding image.

Once synthetic images are generated, a segmentation network (e.g U-net [21]) is trained. This network learns the relation between the synthetic image and ground truth pair to infer image to object relationships (Figure 2a). We evaluated the usefulness of synthetic data for this purpose by applying it to the DeLTA implementation of U-net [22], a popular and robust pipeline for analysing images of cells in linear colonies. Because the output from the U-net is only a probabilistic mask, a correct probability threshold must be determined to compute accurate binary masks. To calculate the optimal probabilistic threshold value, we developed a validation mechanism which utilises synthetic data to estimate the optimal mask pixel probability threshold by passing an independent set of synthetic validation data through the trained model and thresholding the output until the masks are maximally similar to the ground truth masks (2b). Two metrics can be used to evaluate the optimal mask threshold, one is the Jaccard index, which is a pixelwise measure of the difference between the true masks and the target thresholded masks. The other measure involves maximising the cumulative intersection between the length and width distributions of cells in the true masks with those in the target masks. We show that these independent metrics produce the same optimal mask threshold (Figure 2c). Since SyMBac uses synthetic data for training, validation data is not available in the traditional sense. Instead, the model is evaluated automatically epoch-by-epoch in terms of cell identification errors, and the model with the lowest error is kept (Figure 2b and 2e). Automatic identification error rate calculations are possible without validation data due to the predictable nature of bacterial growth in the mother machine (Supplementary Information 7). In general, 100 epochs were enough to drive the identification error below 0.5%.

**Figure 2:**
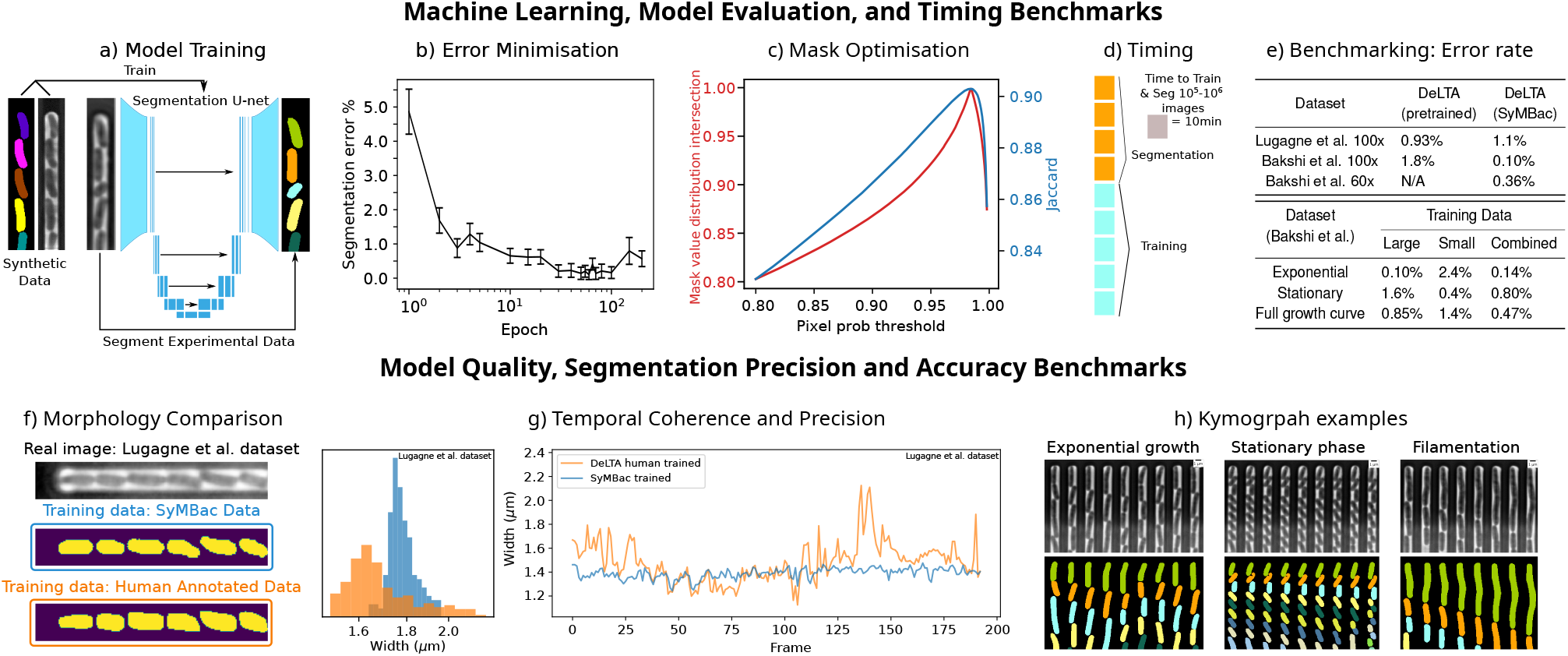
Image segmentation with SyMBac: **a)** Schematic of the U-net model being trained using synthetic data and then segmenting real data to produce accurate masks. **b)** Segmentation error of trained models are evaluated automatically epoch-by-epoch by analysing the sawtooth curves of cell length changes over time (Supplementary Information 7), and the model with the lowest error is kept. **c)** The optimal probability threshold is identified by passing synthetic validation data through the model, and the threshold is adjusted to maximise the Jaccard index between the predicted masks and the ground truth masks, or the maximal distribution intersection between size distributions. **d)** A typical time to train the network (on 2000 images) and segment approximately one million images (Nvidia GeForce 1080Ti). **e)** Top: Performance (cell identification error) benchmarking of DelTA trained with SyMBac. Models trained with SyMBac perform well for both oil and air objectives and outperform models trained with user-generated training data. Bottom: SyMBac enables generation of complete training data to tackle datasets with varying cell sizes, such as along the growth-curve. To optimise the performance throughput the entire dataset, models trained with a combination of large and small cells resulted in best performance across the entire growth curve, while specialised models performed best for the data type they were trained on. **f)** The masks from SyMBac trained models are truer to the geometry of the cells, displaying no aberration when compared to model outputs trained on human annotated data. **g)** The masks also maintain a narrow distribution of widths, while the masks from DeLTA trained with human annotated data display wide variation with the peak shifted to lower values and show 2.5x higher variation in cell width. **h)** Example kympgraphs of 100x data showing cells in a variety of states (exponential growth, stationary phase, filamenting) with accompanying masks, highlighting the robustness of the model trained on mixed data to segment cells of multiple cell sizes and morphologies.

SyMBac synthetic data is accompanied by perfect ground truth masks. Therefore SyMbac trained models produce more consistent, less flawed, and visually more “natural” cell mask shapes compared to models trained with user annotated data (Figure 2f and 2g). This is because masks from synthetic data are never corrupted by changes in the images from which they come. To perform a rigorous comparison between synthetic data and human generated data, we generated a synthetic dataset based on reference images from the original DeLTA paper’s dataset. We then trained two models, one on our synthetic data, and one on the training data provided in the DeLTA paper [22]. In the the DeLTA paper, a semi-automatic tool was used to generate training data, which involves subjectivity in the choice of thresholding parameters and morphological operators. This results in training data with oddly shaped masks, which are consequently learned and then reproduced by the network during training and prediction (Figure 2f). The network trained with SyMBac produces far more realistic mask predictions compared with models trained with human curated training data (dataset from Lugagne et al. [22]) while maintaining almost the same accuracy (Figure 2e). Furthermore, we observe that the DeLTA model trained with SyMBac shows much higher temporal coherence when compared to models trained on human annotated data. Images from the original DeLTA paper, when segmented with the DeLTA model trained with the supplied training data, showed unrealistic fluctuations in cell width from frame to frame, while SyMBac trained models did not (Figure 2f). This manifests as a narrower cell width distribution from SyMBac trained models over the entire experiment (Figure 2g). This improves segmentation precision and makes models robust to large variations in cell size (Figure 2h, Supplementary Information 9), while maintaining the ability to detect very small variations in size due to the segmentation precision. This opens up possibilities of deeper investigations into cell size and shape regulation, as cell width can now be accurately characterised.

DeLTA models trained with SyMBac data were able to achieve comparable identification accuracy (table in Figure 2e as the model trained on human annotated and curated data, which is remarkable, since it shows that SyMBac is able to generate realistic training data from just the reference images. Of course, the true advantages of SyMBac is in its ability to generalise to experimental variations where pretrained models fail. When we tested against a data set collected with a 60x air objective, the pretrained DeLTA model could not produce any masks and therefore we could not quantify the error. Conversely, SyMBac can generate similar images and ground truth pairs, yielding highly accurate segmentation with an error rate of 0.36% (Figure 2e) and high-quality cell masks (examples in Figure 2h and Figure 9). DeLTA trained with SyMBac data also gave 18x better identification accuracy compared to pretrained DeLTA models for a separate 100x dataset (experiments detailed in Bakshi et al. [3]).

Segmentation algorithms (both ML and non-ML) face an even harder challenge when the data type itself changes during the experiment, for example cells changing size, shape, and intensity along a growth-curve experiment as they transition from exponential to stationary phase and back (Figure 3a) [3]. Since cells in the stationary phase are very small, the relative annotation error grows (as shown in our benchmarking results in Figure SI 1) and compromises the performance of the ML algorithm (Figure 1g). On the other hand, non ML methods suffer from changes in object size, as algorithmic parameters need to be retuned when large changes are observed in the image (for example, morphological operators need to have their window size changed to account for different sized cells). SyMBac can address this issue by generating high-quality training datasets for cells at any size (Figure 1e), producing accurate cell masks throughout the experiment (Figure 2h and Figure SI 17). While models trained with small cell training data perform well in stationary phase, this comes at the cost of reduced accuracy in exponential phase, the converse is also true (table in Figure 2i). Instead, a complete dataset, which combines synthetic images of both types, leads to an overall better performance in identification accuracy and highly precise masks for estimating cell length and width (Figure 2h).

**Figure 3:**
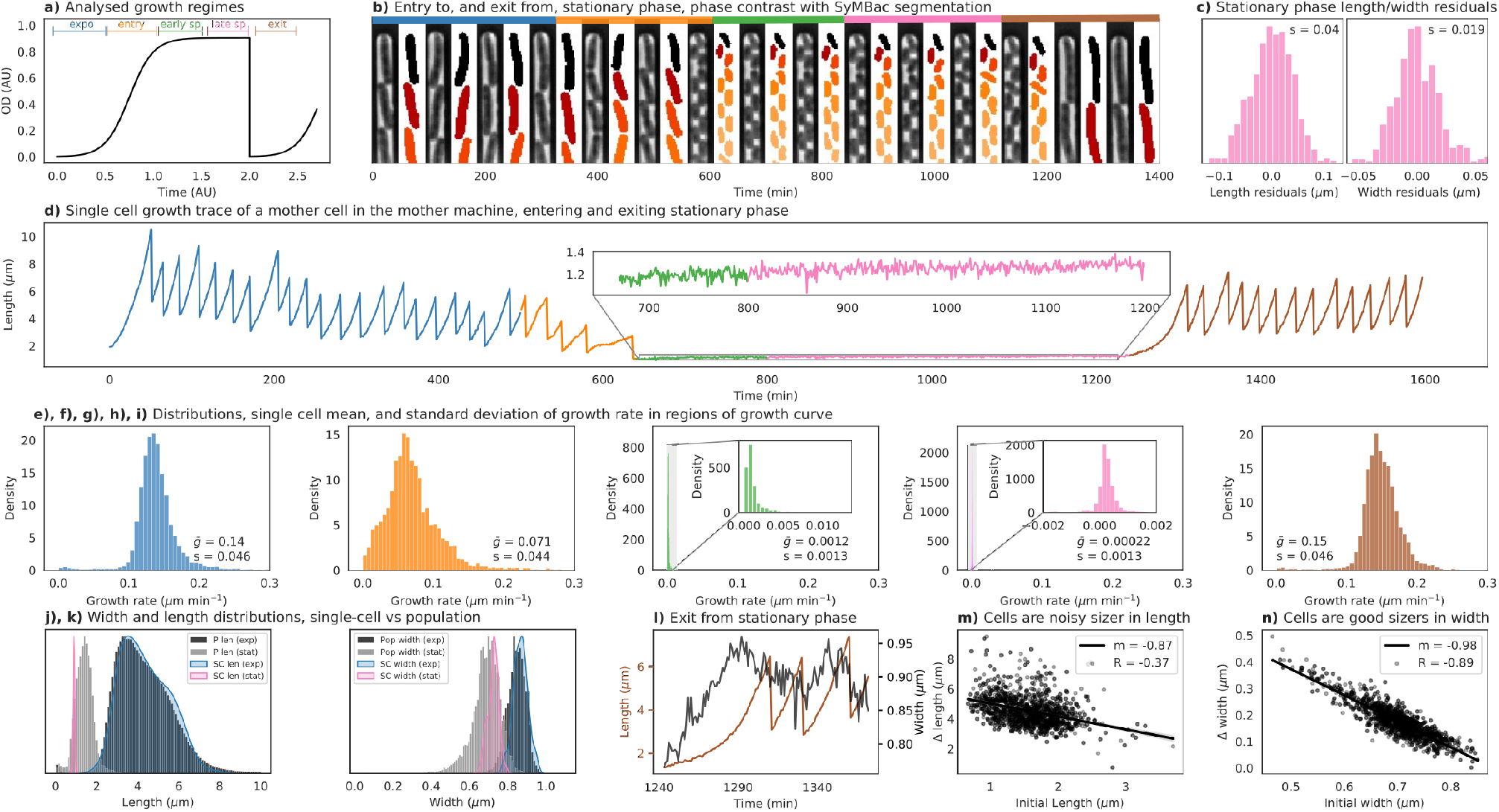
**a)** DelTA trained with SyMBac was used to segment single cell data throughout all growth curve regimes (colour-coded and used throughout the figure). **b)** Sample kymograph with masks of a single mother machine trench going through the growth curve. **c)** The SyMBac trained model produces masks with precisions of 40 nm for length and 19 nm for width. **d)** Example time series of the size of a single cell going through an entire growth curve. Inset shows cell length changes during stationary phase. **e), f), g), h), i)** During the exponential phase, cells exponentially increase their size with a mean growth rate of 0.14 μm min^-1^ which is equivalent to a population doubling time of 19.8 min, consistent with the bulk growth measurements of cells in this rich defined media [3]. The distribution of growth rates shifts to the left as cells enter stationary phase (orange and green phase) and eventually stops 6 hours into the stationary phase (pink). **j) k)** Comparison of the length and width distribution of the population and for an example cell over time. Histograms represent population distributions, whereas filled in curves represent distributions for a single cell. During exponential phase, a single cells will sample the entire distribution of widths and lengths (shown by the blue filled in curve, overlayed over the black population histograms), but upon entering stationary phase, a single cell is locked into specific values of length and width (as shown by the sharp pink curves, overlayed over the grey population histograms). **l)** An example cell trace during exit from the stationary phase, showing the cell’s width and length increase in the wake-up phase. **m)** Comparison of initial length and the added length before the first division after exit from stationary phase shows that cells are noisy as sizers towards length regulation. **n)** Comparison of the initial width and the added width before the division shows that the cell is an almost perfect width-sizer, dividing only when individual cells reach a critical width.

As a proof-of-principle application, we have used SyMBac to analyse a timelapse experiment of cells growing in linear colonies under changing growth conditions, whereby the cells undergo a feast famine cycle [3]. The precise estimate of cell sizes from the masks enables the computation of single cell growth rates during exponential phase, along the entry and exit from stationary phase along a growth curve (Figure 3d), and at different points within stationary phase (Figure 3e-i). During exponential growth, cells grow at a mean rate of 0.14 μm/min, however upon entry to stationary phase, the cell growth rate dramatically reduces by 2-fold, to a population mean of 0.071 μm/min and then asymptotically decreases as cells progress deeper into the stationary phase. However a small residual, but detectable growth rate is maintained (0.000 22 μm/min), which is approximately 640 times slower than the exponential growth rate. This small growth rate can be readily detected due to the higher precision of our segmentation model. This can be seen in Figure 3f-h. The cells then recover their growth rates as they enter the exponential growth rate when fresh media is reintroduced (Figure 3i). Since the cells almost stop growing in the deep stationary phase, the fluctuations in the cell size estimates from the binary mask can be used to estimate the precision of segmentation (Figure SI 14). We estimated the precision of the output from SyMBac models to be 40 nm along the length and 19 nm along the width (Figure 3c). The highly precise estimates of cell size and shape enable us to look at shape regulation at different stages of starvation and resuscitation.

Most interestingly, however, SyMBac trained models allowed us to very precisely quantify single cell width changes. During entry to stationary phase, the length and width of individual cells drop during division and settles heterogeneously at various stages as the division halts. This leads to highly variable cell sizes in the stationary phase across the population. This is demonstrated by the large population variance compared to single cell fluctuations in length and width (Figure 3j, k). It can be seen that while in exponential growth, a single cell will sample the entire population’s width and length distribution, however upon entry to stationary phase a single cell is “locked” into a fixed width and length. During resuscitation, cells increase their width and their length and reach homeostasis during exponential phase, as demonstrated by the comparable variability across the population and individual cells. We also find that, during the wake-up phase, cells divide once they have reached a critical size (Figure 10). Upon further investigation, when we look at the length and width regulation during this phase, we find that cells more strictly regulate their width than their length during wake up and each cell divides upon reaching a critical width (Figure 3m vs Figure 3n). This shows that cells regulate their size through width control, which is reasonable, as the volume of a cell has a stronger dependence on width than length (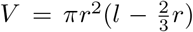). The large variability in cell width in stationary phase coupled with this strict width requirement for initiating cell division leads to a strong correlation between the shape and size of individual cells and their resuscitation behaviour, with important implications for persistence towards antibiotics, and population fitness in general.

Finally, while we demonstrated that SyMBac is most powerful in its ability to train highly accurate models for image segmentation of cells growing in linear colonies, we also sought to extend SyMBac to produce synthetic micrographs of cells growing in monolayer colonies, such as the “biopixel” device [23], 2D turbidostat [16], and even colonies growing on agar pads. For this task we replaced SyMBac’s cell simulator backend with CellModeller [24], a powerful cell simulator, more suited to 2D growth than our custom simulator. We then added physical constraints to our model to simulate the walls of a microfluidic turbedostat, like the one described in [16] (data generously provided by the Elf Lab, Uppsala Univeristy). We also simulated growth of micro-colonies on agar pads in both fluorescence and phase contrast. SyMBac was able to simulate images very similar to real micrographs (Figure 4). Modifications to the simulation, the point spread function and the segmentation pipeline are described in Supplementary Information 11. These synthetic images and the corresponding ground truth positions were used to train deep-learning networks to achieve good segmentation of experimental images from these platforms and produced masks of unprecedented qualities, in terms of the shape of individual cells and robustness towards the variation in size and contrast.

**Figure 4:**
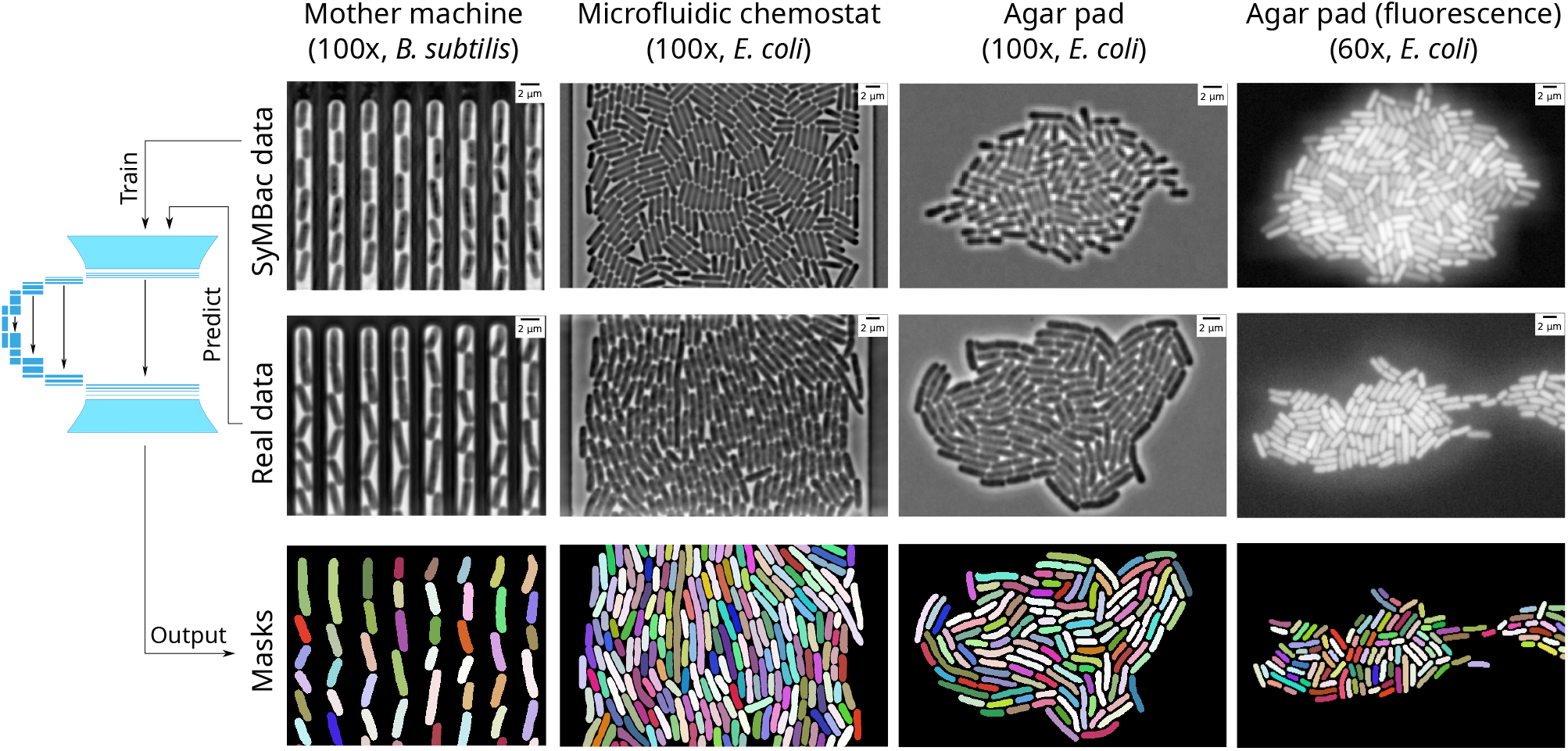
Extensions of SyMBac for cell segmentation in images of linear colonies of *B. subtilis* (very straight cells, unlike *E. coli* which have more curvature), monolayers colonies in a 2D microfluidic turbidostat chamber (data from [25], kindly provided by the Elf Lab, Uppsala University), growing colonies on agar pad, and low resolution fluorescence snapshots of dense cell clusters on agar.

## Discussion

Here we have demonstrated a new approach of cell segmentation using deep-learning models trained entirely on synthetic training data. Synthetic data has been used previously to aid in segmentation. MiSic [26] is a tool for segmenting bacterial micrographs based on real training data, but supplements this data with “synthetic” data, which however does not accurately reflect the optics, physics, or biology of the experiment which is being analysed, and thus cannot segment images based purely on synthetic data. BCM3D [27] is a tool used for fluorescence biofilm segmentation, however it relies on deconvolution, which can lead to increased noise and phantom objects in the output masks, or experimentally derived PSFs. Additionally it is not compatible with linear and monolayer colonies that are typically used in high-throughput quantitative experiments, nor is it compatible with phase contrast images, which are ubiquitous in the imaging of bacteria. SyMBac is the first tool which can generate synthetic data by modelling the entire experimental setup, end-to-end, and is compatible with phase-contrast and fluorescence images from various imaging platforms, such as microfluidic devices and agar-pads.

Training data generated in more traditional ways, using various thresholding algorithms results in variable masks for cells of identical shape and size when the cell’s pixels have different brightnesses, illumination profiles, etc (the same cell would be assigned a different training mask depending on the preprocessing algorithms and parameters used). This introduces uncertainties in the true pixels for each cell, which is passed on to the model from training data, and is propagated to the output. At the same time, while input training masks need preprocessing, so do the output masks [22, 12, 13]. Post-processing these outputs will deform the already artifactual cell masks even more, and make cell morphology analysis highly susceptible to error (both systematic and random). Since SyMBac data is accompanied by perfect ground truth information, the resultant masks are highly accurate and precise, and don’t need any post-processing. This eliminates user subjectivity in data creation, curation, and final analysis, increasing reproducibility. The highly accurate masks from models trained with SyMBac have revealed novel insights into cellular growth and physiology under changing conditions and paves the way for systems-level analysis of the underlying factors using such high throughput data.

In summary, SyMBac provides a major technical advancement for the easy adoption and robust implementation of machine learning approaches for analysing microscopy data of microbes. Through rapid and easy generation of high quality synthetic data with large coverage of experimental setups and cellular morphologies, SyMBac addresses a critical obstacle in the application of ML tools for high-throughput single-cell image analysis from various platforms and organisms. Beyond the compatibility towards variations in cell morphology and imaging platform designs, the microscope model in the pipeline is compatible with objectives of various resolutions, including low-resolution air objectives. Therefore, SyMBac enables easy creation of training data for low-resolution microscopy images, which is near-impossible for human annotators or contrast based computational programs. This enables high-accuracy segmentation from images collected with air objectives (Figure SI 18), which will have important implications in analysing high-throughput image-based screening experiments ([28, 3, 29, 30]), where low resolution air objectives provide the necessary travel distance, speed, and large fields of view. As SyMBac enables easy creation of synthetic training data at any spatiotemporal resolution, it also addresses the robustness and reproducibility problems with benchmarking machine-learning (ML) and non-ML approaches for cell segmentation and tracking. Therefore, we believe SyMBac will play a critical role in improving performance of the ML algorithms, while also enabling accurate assessments of their performance limits.

## Methods

### Image Generation

Our method employs several steps which make up a pipeline:

- Simulation of growth and division of cells and their interactions with each other and the microfluidic device.
- Extracting geometric information and rendering images of the bacteria and device in the scene.
- Microscope model for generating phase-contrast and fluorescence versions of these images.
- Image optimisation to further improve similarity to experimental images.
- Training ML segmentation algorithms with synthetic images.
- Performance testing.
- Model evaluation.

We give brief descriptions for each of these steps below, with additional information presented in the supplementary material.

### Simulation of growth and division of cells and their interactions with each other and the microfluidic device

We built an agent based model (ABM) of cell growth and division, taking into account the three size regulation modes of bacteria (adder, sizer, timer), while also adding variability to the key parameters governing cell division. Cells are agents with properties such as length, width, position, age, and can be customised to include new properties such as time-dependent fluorescence intensity. These cell agents exist as dynamic objects in a rigid body physics world called a “scene”, modelled using Python bindings of the popular physics engine Chipmunk (pymunk) [31]. We add static objects to these scenes which are shaped into a mother machine trench geometry, with configurable length and width. A global event loop keeps track of cell growth, divisions, inter-cellular collisions and cell-trench collisions at each time-step. A simple API was created to allow simulations to be run by entering the desired distributions of cell lengths and widths by defining the mean, variance and size-regulation mode, and then selecting the desired trench geometry (Supplementary Information 2, Figure SI 10). The simulation can also be watched in real time to check that the chosen parameters are valid (Figure SI 5).

### Extracting geometric information and rendering images of the bacteria and device in the scene

After simulations are complete, geometric information about the cell and trench positions are extracted. This includes each cell’s vertices in every time-point. The cell’s 3D structure is a spherocylinder, which affects how light interacts with it, yet the simulation engine runs only in 2D. Therefore to render 3D cells, the cell’s vertices are revolved along their major axis to generate each cell’s spherocylindrical hull. All of these 3D cells are then morphed to simulate cell bending and then re-rendered and in a 3D scene, computationally represented by a 3D NumPy array (Figure SI 7). A 2D projection of this 3D array is then taken, such that an image is generated where the intensity in each pixel is proportional to the optical path length (OPL) in that region (Figure SI 6). The details of this calculation are provided in Supplementary Information 3. This simulated image corresponds to what would be seen if a microscope had no PSF, and thus induced no diffraction artefacts in an image. From the simulated images at this stage, we also save the ground truth images of the cell positions, which act as the masks in our training data.

### Microscope model for generating phase-contrast and fluorescence images

The microscope model generates images from the scenes by convolving them with PSFs of relevant optics. First, the raw image is rendered at a high resolution (typically 3x the real pixel size) and then convolved with a variety of fluorescence or phase contrast PSFs (also rendered at higher resolution) depending on the application. After PSF convolution, the image is sub-sampled back to the original resolution. Convolution at higher resolution is more accurate, because sub-resolution features of the PSF (such as the concentric rings) are maintained, and propagated to the image. Otherwise, the PSF convolution results would be corrupted by low resolution artefacts. The user can select an ideal fluorescence PSF or phase contrast PSF which are modelled as an Airy disk and obscured Airy disk [32] respectively (Equations 2 and 3 in Supplementary Information 4). The PSFs needs to be parameterised by inputting the microscope objective’s numerical aperture, refractive index of the medium, emission wavelength, camera pixel size, and phase ring and annulus dimensions (details provided in Supplementary Information 4). Upon defining the PSF of choice, it is convolved over the raw image to produce a synthetic micrograph.

### Image Optimisation

Next, the user can use the optional image optimisation to maximise the similarity between the synthetic image and an example real microscope image. Multiple image manipulations are possible, including intensity histogram matching, noise matching, rotational Fourier spectrum matching, camera noise simulation, stage drift simulation, and even manual intensity corrections. This is done using an interactive IPython Notebook interface, where the user is presented with a side-by-side comparison of a synthetic image and a real image from their experiment. Two plots showing the relative errors in intensity and variance between cells, the mother machine device, and the space between cells are given so that the user can manipulate their image to minimise this error (Supplementary Information 5 and Figure SI 10). (Note: While examples of black box optimisers for image similarity matching are included, we avoid their use for this step due to the very noisy error-landscape between synthetic and real images. Moreover, we were unable to define a perfect objective function which guarantees perfect image similarity). Synthetic images are then generated according to the optimal parameters. These parameters can be varied according to a uniform or Gaussian distribution during the image generation process to simulate minor fluctuations in the experimental setup. This also acts as a form of data augmentation, but occurs during the image generation process, which preserves the mapping between object and image.

### Testing

Interfacing code for training machine learning models using DeLTA [22] and StarDist [33, 34] are provided. For most analyses, U-net models were trained with DeLTA, as this provided the most accurate masks. StarDist, while not pixel-perfect, can be useful for especially low magnification data (Supplementary Information 6) because of its robust ability to separate densely packed star-convex shapes.

For mother machine experiments, we tested SyMBac against 4 datasets, which have their details given in Table 1. 4 sets of data were generated, each with approximately 2000 synthetic images. 3 of the datasets are images of *E. coli* growing in a mother machine, taken with a 100x oil objective, and one dataset was imaged with a 40x air objective, using a 1.5x post-magnification lens, giving an effective magnification of 60x. However synthetic images for this dataset were still generated according to the optics of the 40x objective, and then scaled to the appropriate pixel size.

**Table 1:** Datasets analysed using SyMBac

| Dataset | Optics | Biology | Notable features |
| --- | --- | --- | --- |
| Lugagne et al.<br>(exponential) | 100x oil | *E. coli*<br>in balanced growth | Poor contrast data |
| Bakshi et al.<br>(exponential) | 100x oil (NA = 1.30) | *E. coli*<br>in balanced growth | High contrast data |
| Bakshi et al.<br>(stationary) | 100x oil (NA = 1.30) | *E. coli*<br>in stationary phase | Small stacked cells |
| Bakshi et al. 60x<br>(exponential) | 40x air + 1.5x post mag<br>(NA = 0.95) | *E. coli*<br>in balanced growth | Low resolution +<br>non-uniform illumination |

### Model Evaluation

For evaluation purposes, the DeLTA implementation of U-net was used to train models using synthetic data. Because the data is purely synthetic, validation data in the traditional sense does not exist. For this reason we saved the model every epoch and evaluated performance through the analysis of growth data. Bacterial growth in the mother machine traces out a predictable sawtooth wave (Figure SI 11). These sawtooth waves and their derivatives were analysed for spurious peaks, which are the signature of over and under-segmentation errors (Figure SI 12). Detailed discussion is included in Supplementary Information 7.

The output from a U-net is a probabilistic threshold image. This image needs to be converted into a binary mask by thresholding pixel values between 0 and 1. Thresholds close to 0 will generate larger, connected masks, and thresholds close to 1 will generate smaller less connected masks. In order to identify the optimal threshold value which generates the most accurate masks, an independent set of synthetic data is segmented using the neural network, and the probability thresholds are adjusted to maximise the Jaccard index between the ground truth and the model output (Figure 2c). Depending on the type of dataset this value can range between 0.6 and 0.99. If the thresholding value is low (around 0.6), then masks will be connected. To alleviate this we use seeded watershed segmentation to cut masks appropriately. Seeds are generated by first thresholding masks at very high probabilities (*P* > 0.999), and performing the watershed on the optimal probability binary mask. It must be noted that we do not perform any post-processing on the training data or the segmentation results which would affect the shape of the mask. After generation of the masks, information on cell curvature, radius (width) and length are calculated using the Colicoords Python library [35] if further analysis is desired.

### Extension to 2D growth

We have extended the use of SyMBac beyond experiments with linear colonies, to include growing monolayer colonies in microfluidic devices and agar pads. For this purpose the cell simulator back-end was switched from our custom implementation (which is optimised for 1D growth in linear colonies) to CellModeller [24], a more general cell simulator for 2D colonies. Scenes are redrawn in the same way as previously described, and either phase contrast or fluorescence PSFs are convolved to generate respective image types. The main difference here is the addition of additional phase objects to simulate other microfluidic geometries (shown in Figure 4), and pseudorandom (Perlin [36]) noise to simulate the textures seen in typical phase-contrast images of microcolonies on agar pads (Figure SI 22). Additionally, the point spread function was adjusted by adding a very small constant offset to modulate the amount of shade-off and halo effects which are characteristic of phase contrast images (Figure SI 21). The details of this process are described in Supplementary Information 11. For 2D geometries, 20 CellModeller simulations were run starting from a single cell, generating in total approximately 1000 unique cell images. This was further augmented by sampling a large variety of parameters, which ensures training samples have variable background noises, shade-off and halo amounts, and cell positions. This can be considered a form of data augmentation, but this step is performed before training the model, rather than during, and is based on the type of image one wants to segment. In total each model is trained on 5000 unique synthetic images.

## Code availability

SyMBac is available as open source software, released under the GPL-2.0 licence.

Code is available at https://github.com/georgeoshardo/SyMBac

Additionally, for those who do not wish to use an API through interactive notebooks, we are committed to providing a GUI for SyMBac in due course.

## Acknowledgements

We thank Hannah Earley and Kavi Shah for very detailed feedback on the manuscript.

We thank the Dunlop Lab for producing the easy to use DeLTA, and for their permission to include the source code in SyMBac’s codebase.

We thank the Paulsson Lab, Harvard Univeristy, for providing the agar pad and mother machine data data, which was captured by S.B. We thank the Elf Lab, Uppsala University, for providing the microfluidic turbedostat data.

This research in the S.B.’s laboratory was supported by the Wellcome Trust Award [grant number RG89305]; a University Startup Award for Lectureship in Synthetic Biology [grant number NKXY ISSF3/46]; Royal Society Research Grant Award (Award number G109931); and the Ph.D. Studentship from BBSRC (to Georgeos Hardo)].

## Author Contributions

G.H: Conceptualisation, Methodology, Software, Experiments, Visualisation, Analysis, Writing. M.N: Conceptualisation, Methodology, Software, Pilot and Feasibility Study. S.B: Conceptualisation, Experiments, Analysis, Writing.

## Supplementary Information

### 1 Benchmarking performances of traditional methods of generating cell masks

To compare the accuracy of other segmentation methods, such as human annotated training data, traditional Otsu thresholding, and cell perimeter evaluation using membrane dyes, we generated synthetic images of single cells in phase contrast and with membrane dyes, along with accompanying ground truth. We then segmented the cells with Otsu’s method and the membrane dye method, comparing the output masks to the ground truth. We show in this section that these methods systematically underestimate the cell’s dimensions.

#### 1.1 Human drawn masks

In order to test how accurate humans are at annotating images to generate training data, we sent 3 researchers a set of 100 synthetic images. The researchers were asked to both label the images and time themselves. The labelling was performed by sending each researcher a Python script which would open a napari window [1], and allow them to manually segment cells, saving their output to a file. We then compared the length, width, and pixelwise (IoU) accuracy of the human generated masks to the ground truth masks of the synthetic data. The IoU output of this result is given in Figure 1i, showing that humans consistently perform poorly in segmenting cells, especially if they are small and in stationary phase. We also observed that there was a significant bias in the way in which humans were mis-segmenting data, based on the cell’s dimension and growth phase, shown below in Figure SI 1

**Figure SI 1:**
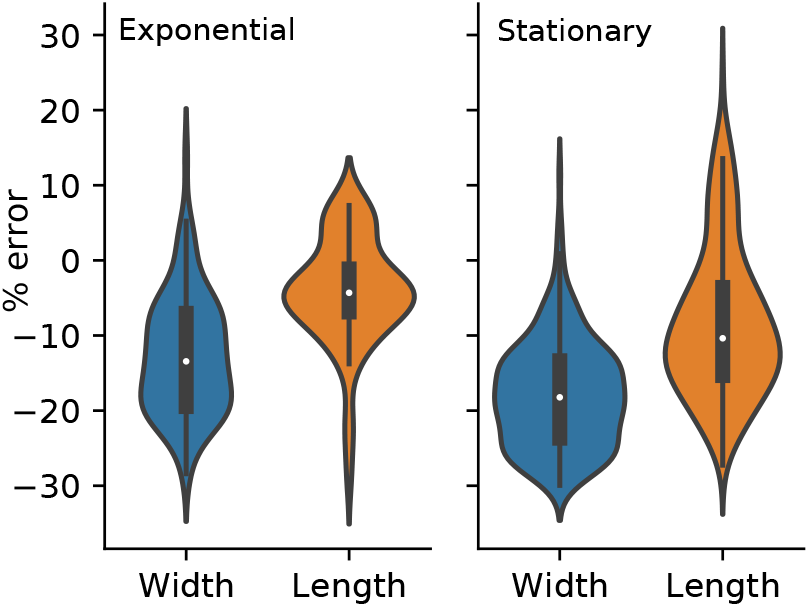
The human annotation error is quantified by 3 researchers who were asked to manually segment 100 synthetic images. Their segmentations were compared against the ground truth. The worst performance was always in the width dimension with this becoming worse when the cells were in stationary phase. There was also a systematic tendency for human segmentations to underestimate the cell’s dimensions on average.

#### 1.2 Membrane Dyes

Next we show that using membrane dyes for estimation of cell size using diffraction limited imaging leads to severe underestimation of cell perimeter. We simulated images of cells tagged with a membrane dye by modelling cells as spherocylinders. We then assigned high fluorescence intensity values to the hull of the cell and performed 3D convolution of a fluorescence PSF approximated as a 3D Gaussian with appropriate X,Y and Z parameters. We then compared the true width of the cell with what one would infer if they assumed the cell width was the inter-peak distance of the maximal fluorescence intensities across the cell’s minor axis. The simulations and a sample of a cell are shown below in Figure SI 2.

**Figure SI 2:**
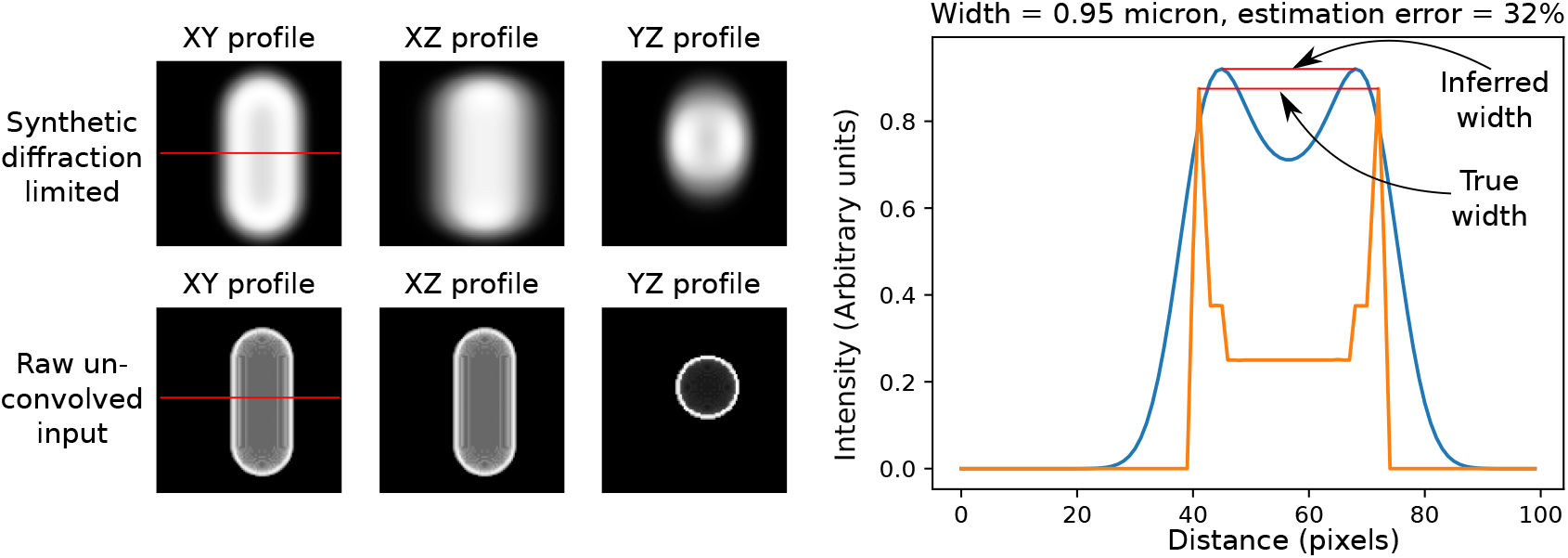
**Left:** Simulation of cells tagged with membrane dyes. Red line shows the slice taken for the intensity profile in the XY plane. Membrane dyed cells are simulated as hollow spherocylindrical hulls, with 0 intensity in the cytoplasm, and an intensity of 1 in the membrane. 3D convolution is then done with a Gaussian approximation to the fluorescence PSF to simulate the raw microscope image. **Right:** Sample intensity trace of the intensity across the width of a cell. Blue shows the intensity of a diffraction limited image of a membrane dye tagged cell, and orange shows the true intensity profile. The microscope’s optics corrupt the image and lead to a decrease in the inter-peak distance which subsequently underestimates the cell width.

This error becomes increasingly large as as the cell’s width gets ever smaller. Figure SI 3 shows the error rates for width estimations for simulated cells from 0.6 microns to 1.4 microns in width. A very narrow cell, with a width of 0.6 microns would have a width estimation error of close to 80%. This represents a best case scenario, as the images were convolved with an ideal point spread function, and no noise was added to the image. In reality the estimation error will be worse.

**Figure SI 3:**
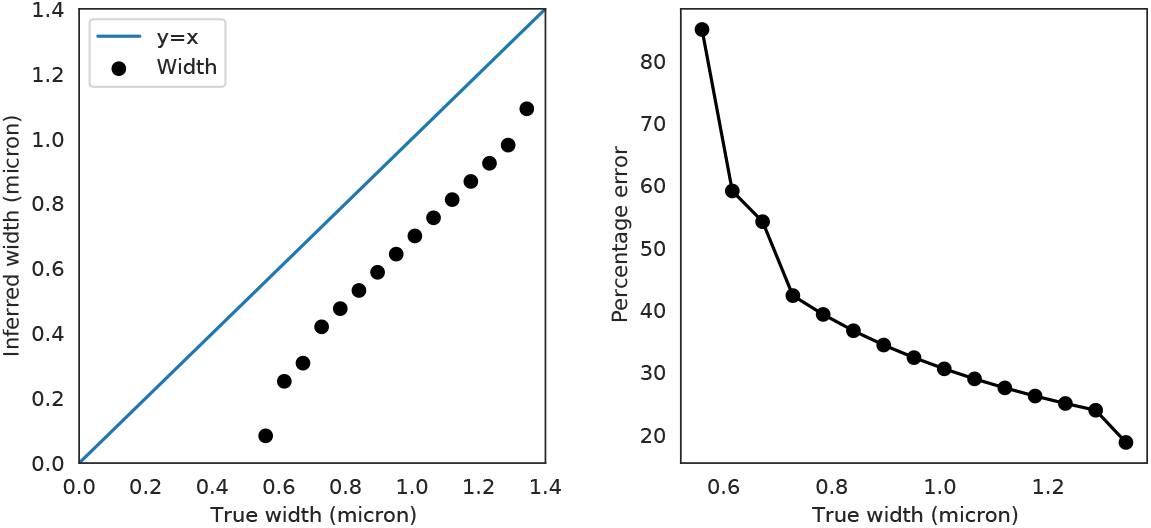
Left: there is a systematic error in measuring the cell width using membrane dyes. Width is consistently underestimated. Right: The error in width estimation by using the inter-peak distance of membrane dye intensity is very high. A typical cell with a width of 1 micron will suffer an estimation error of more than 30%. Additionally for very thin cells it becomes impossible to make out the perimeter due to reaching the diffraction limit.

If membrane dyes are the only way to measure width due to experimental constraints (e.g the lack of a phase contrast objective), then packages such as ColiCoords [2] are recommended, as they can mitigate the error incurred by this method by allowing the user to choose the method of perimeter estimation.

#### 1.3 Otsu’s Method

While Otsu’s method alone rarely yields good results for the segmentation of phase contrast cells, it is often used in combination with other preprocessing steps in the generation of training data for further use in machine learning pipelines. Thus its error must be investigated, because if it is used in the training data generation pipeline, its error will be propagated to the learning algorithm chosen. We simulated single cells imaged under phase contrast optics in order to quantify the error from this thresholding method compared to the ground truth mask. The error is systematic and constant across cell widths, meaning that narrower cells suffer from greater relative error rates in dimension estimation (Figure SI 4).

**Figure SI 4:**
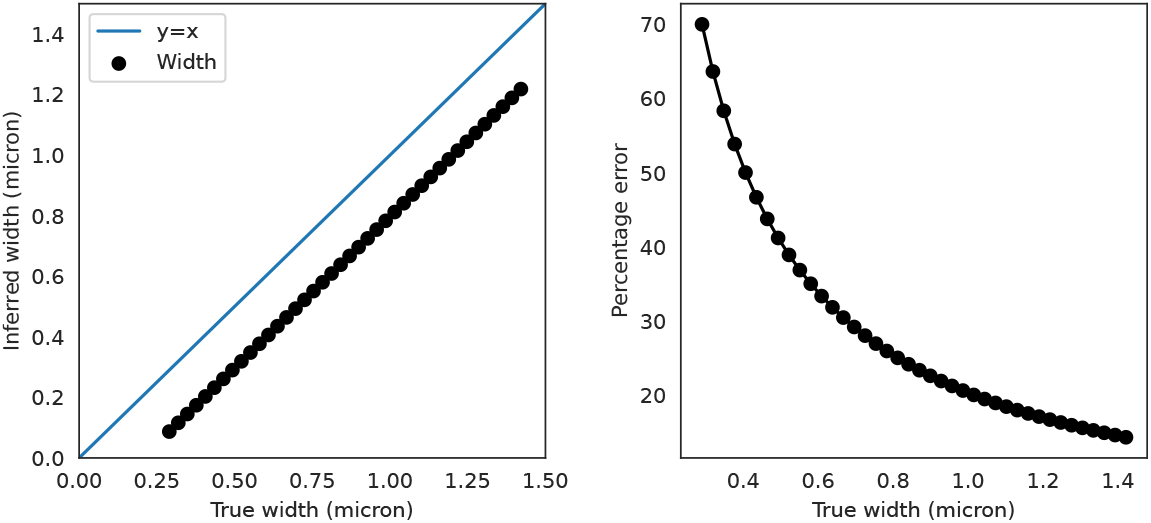
**Left:** The error from Otsu’s method is a consistent offset for all cell sizes. While one could in theory compensate for this, the offset will not be immediately calculable as each microscope’s PSF will slightly vary. **Right:** the relative percentage error grows quickly as cells become narrow.

These data show that traditional methods of segmenting cells, while fine for identification of the cell, fail to accurately measure a cell’s dimensions. Membrane dyes underestimate the width, while Otsu’s method does the same. These underestimation errors would be propagated to a learning based algorithm if used to generate training data. Further segmentation methods, such as those based around local thresholding and watershed may yield better identification accuracy, but still suffer from errors in dimension estimation. This is because one needs to constantly tweak subjective parameters until the segmentation “looks right”. For bioimage analysis of large objects, such as eukaryotic cells, this is typically not a problem, but as shown, it manifests for small objects which exist at the diffraction limit, such as bacteria.

### 2 Details of Agent Based Model and Rigid Body Physics Simulation

Our cellular simulation is an agent based model, where each cell is defined by a number of attributes (`length, width, resolution, position, angle, space, dt, growth_rate_constant, max length, max_length_var, width, width_var`). These attributes are unique to each cell, except for dt which defines the integration time-step for the cell’s growth. Cells can grow according to the adder, sizer or timer growth model.

For the spatial component of the simulation we used Python bindings for the popular Chipmunk physics engine, called Pymunk [3] in order to create a custom simulation environment. Cells exist as dynamic objects in a Pymunk space and can move around. Each time-step, the lengths of the cells are updated. This causes some cell hulls to overlap, the physics engine is then called to resolve these conflicts and move cells. While simulations most often produce realistic cell-stacking dynamics, the nature of cell-cell and cell-trench interactions can be varied by adjusting the number of physics iterations in each time-step, and by adding gravity to the simulation. Stronger gravity in the direction of the bottom of the trench will induce tighter cell stacking, and in some cases double-loading of cells into the mother machine (which is a common occurance in experiments where the trench is too wide for the organism of choice). Low gravity, or negative gravity results in cells seemingly repelling each other, keeping a large distance from one another in the mother machine (often seen with motile strains which can leave trenches mid-experiment). Due to the unpredictability of cell stacking dynamics in real experiments (which vary due to media flow rate, cell size, trench dimensions, etc) we leave these parameters free to be changed by the user in order to maximise the similarity of the simulation to their experiment. The simulation can be watched in real time while it is running (Figure SI 5).

**Figure SI 5:**
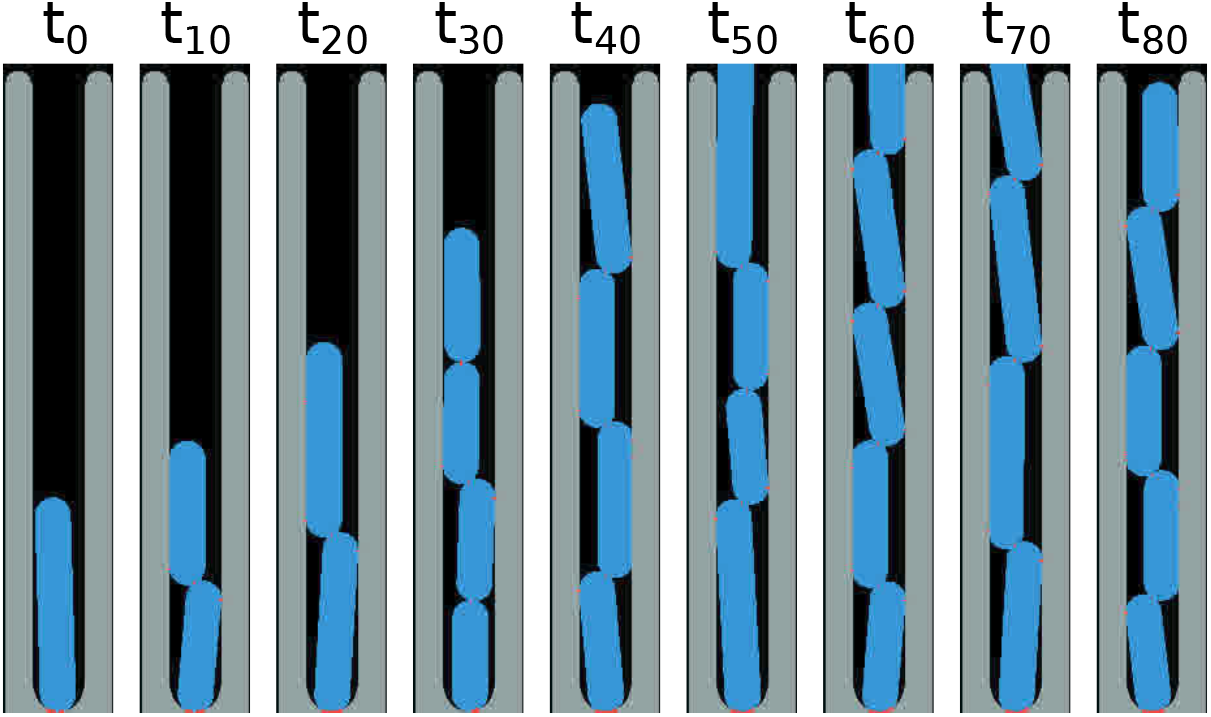
Example kymograph of cells in the trenches from a simulation run. The user can monitor this simulation in real time and adjust parameters to achieve the desired result.

### 3 Details of OPL Calculation and Scene Drawing

At every time-point, each cell’s location and dimensions are kept track of and recorded. This data is used to redraw the entire scene in 3D. While the simulation itself is not performed in 3D, with all interactions restricted to the XY plane, it is important to simulate the optics in 3D. The optical path length (OPL) of each pixel in the image must be simulated correctly. This is the optical distance that a ray of light experiences while travelling through the sample, and is given by a product of the geometric (real) distance travelled, and the refractive index of the medium which the light passes through. Our simulation has 3 main objects which light can pass through: the PDMS of the mother machine, the cell growth medium, and the cell itself (Figure SI 6). This corresponds to 3 refractive indices. PDMS and growth medium have a constant depth in the XY plane, however the cells do not, this is the main reason for the importance of the 3D treatment of the cells. Light has a higher optical path length down the centre-line of a cell than at its edge.

**Figure SI 6:**
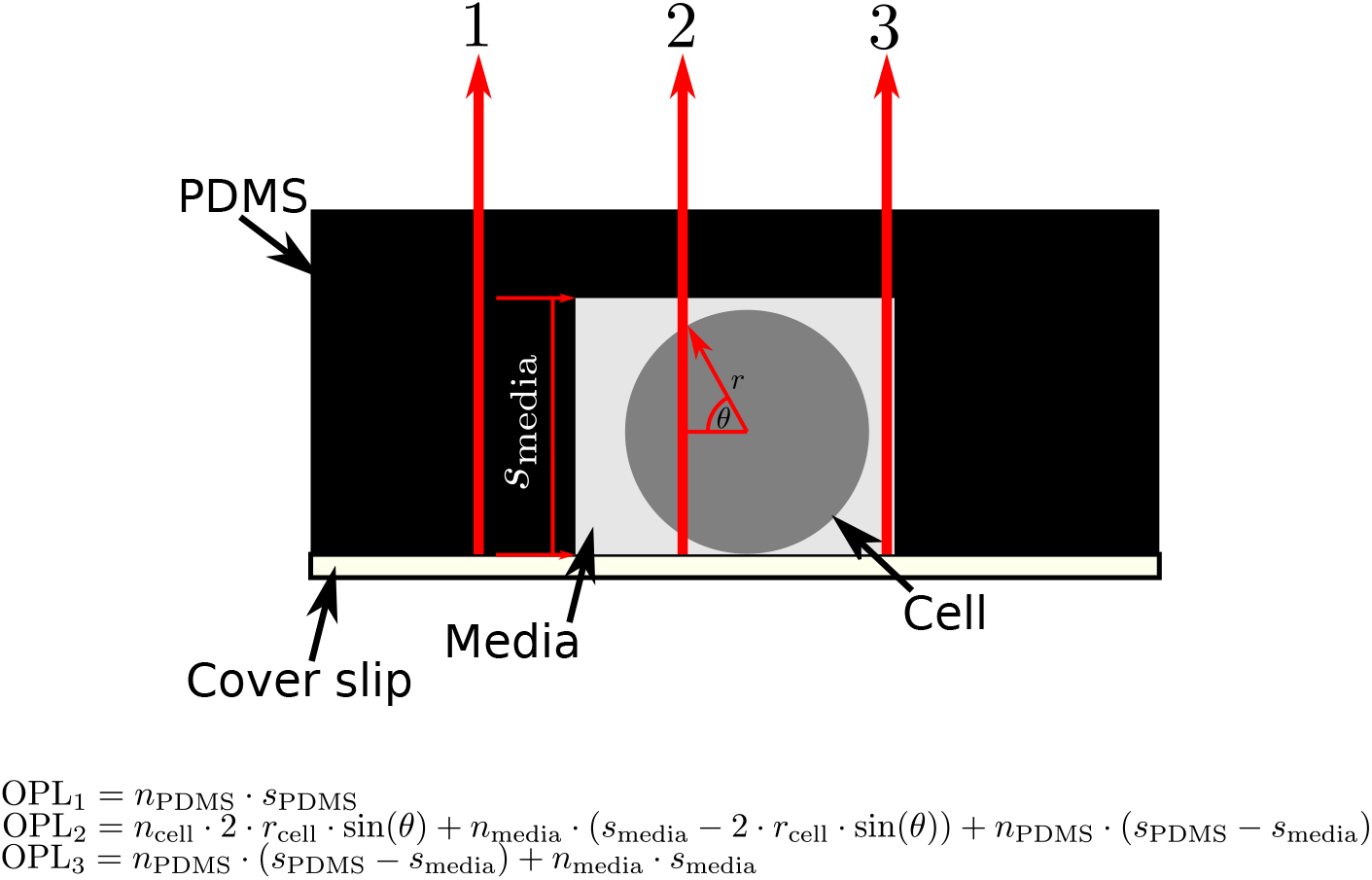
The OPL value is calculated for each pixel according to geometry and relative differences in refractive index, however the exact values of the refractive indices are left free to be optimised during the image generation step.

The 3D OPL images are then optionally modified to create curves in the cells. Each cell, being represented by an array can be morphed. In order to do this we roll individual rows of the cell image by a random number of pixels, with the transformation given by:

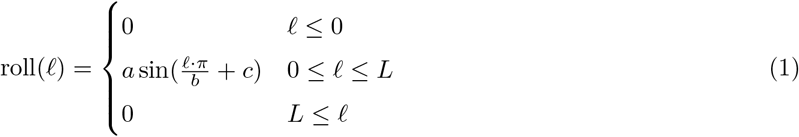

where *L* is the total length of the cell, *l* is the position down the length of the cell, and *a, b,* and *c* are randomly chosen for each transformation, but kept constant for individual cells. An example of the transformation of an OPL image viewed in the XY plane is shown in Figure SI 7

From the 3D OPL, masks are then generated. Two types of masks can be generated: instance masks, where each cell’s mask is given a unique value, but no zero value pixels separate different cells, and semantic masks, where all cell masks have the same value (1), but cells are separated from one another with zero value pixels. Instance masks are intended for use with Star/Splinedist, while semantic masks are for use with DeLTA.

**Figure SI 7:**
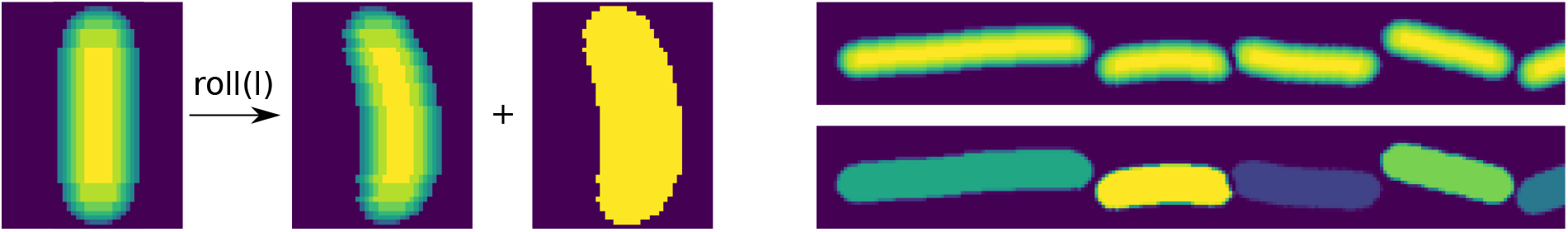
Left: A random transformation of a straight cell (OPL image in XY) into a curved cell allows for rendering of more realistic images of mother machine images. The mask of the cell is then simply taken as the entire region defining the newly transformed cell. Right: Example output from the simulation of a single frame OPL image and its corresponding instance mask output.

After this process is completed, the trench is drawn around the cells according to the simulation input dimensions.

### 4 PSF Definitions and Convolution

The fluorescence point spread function is modelled as a standard Airy disk, given by:

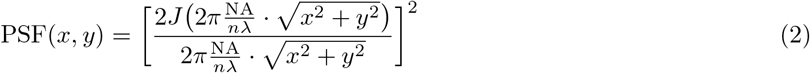

Where *J* is a Bessel function of the first kind, *n* is the refractive index of the imaging medium, NA is the numerical aperture of the objective, *λ* is the emission wavelength, and scale is the pixel-size.

The phase contrast PSF is modelled similarly to [4], as an obscured Airy disk given by

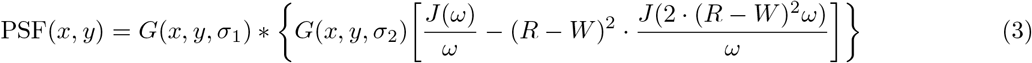

with

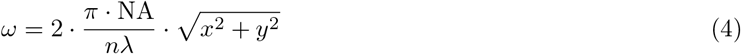

Where *R* and *W* are dimensions of the phase ring and condenser annulus (a diagram of which is shown in Figure SI 8). The phase contrast PSF is also convolved with a gaussian kernel (*G*) to simulate defocus [5], and multiplied by a 2D gaussian to simulate apodisation. Example PSF images can be seen in Figure SI 9.

**Figure SI 8:**
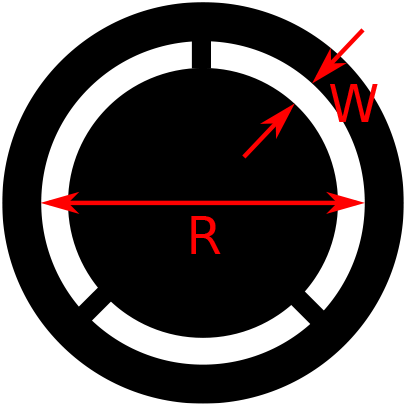
Diagram showing the diemsions of the phase ring and condenser annulus to be used with Equation Equation 3 to parameterise the phase contrast point spread function

**Figure SI 9:**
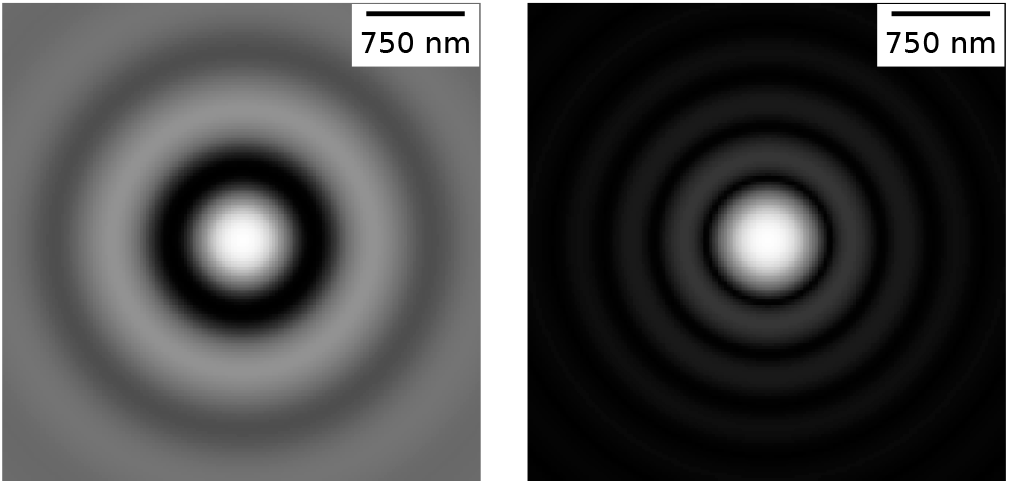
Phase contrast (left) and fluorescence (right) kernels generated with *λ* = 0.75 micron, NA=0.97, *n* = 1, *W* = 0.8 mm, *R* =5 mm. The phase contrast kernel is displayed with no apodisation or defocus. Both kernels have their intensities square rooted to enhance contrast for the reader.

Convolution is done discretely over the OPL images on the GPU using CuPy. OPL images and kernels are rendered at at least 3x the imaging resolution. This ensures that when the convolution takes place, a high resolution kernel is convolved, and no details of the kernel’s concentric rings are lost to pixelation. Only after convolution has taken place is the synthetic image then resized back to the camera’s original pixel size.

### 5 Image Optimisation

After convolution and resizing, optional image optimisation takes place. The first optimisations are tuning of the refractive index of the PDMS, media, and the cell. In order to get the correct intensity values for these features, the users are presented with an interactive Napari [1] window (Figure SI 10). The user must label (with any value > 0) individual layers corresponding to the cell, media, and the device. Accuracy is not needed as only simple estimate of the mean and variance of each intensity is required.

Next, the lens apodisation (modelled as the PSF multiplied by a 2D gaussian) and defocus (modelled as the PSF convolved with a 2D gaussian) can be specified by varying the sigma values in Equation 3. These optimisations alone are often sufficient to produce a somewhat similar image, however similarity can be maximised by the addition of noise and matching various properties to real images. Noise is modelled in two ways: photon shot noise is modelled in each pixel as Poisson noise with mean equal to the pixel’s intensity, and dark noise is modelled in each pixel as the addition or removal of normally distributed intensity with user controlled variance. After the addition of noise, histogram matching between the synthetic images and a representative real image takes place. Histogram matching resolves any remaining discrepancies in image intensity. Additionally, in some instances rotational Fourier spectrum matching can be beneficial, and this is implemented through a python translation translation of the SHINE toolbox [6]. Fourier spectrum matching is only recommended for high resolution and high magnification images, as its main purpose is to replicate the intricate texture found on cells (which is typically not visible on lower magnification images). In our pipeline we only turn on this setting for 100x oil images. All of these parameters can be adjusted interactively in an IPython notebook using sliders, shown below in Figure SI 10.

**Figure SI 10:**
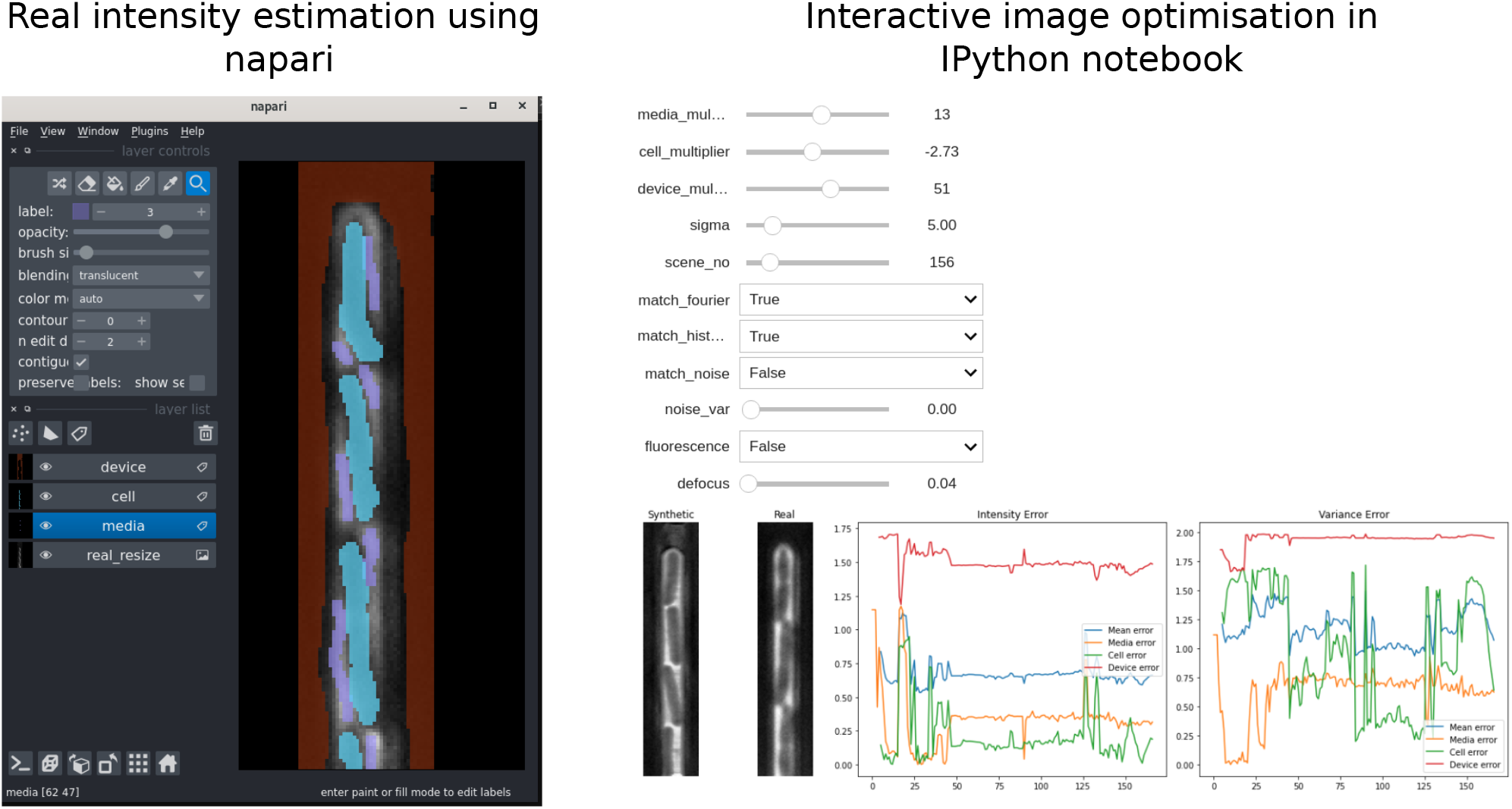
Left: Labelling of a sample real image to identify image regions (cell, device, media) in order to adjust the intensities of the conrresponding regions in the synthetic image. Right: Example of the interactive adjustment interface mid-optimisation for a 40x image. Adjustment sliders for the media, cell and device intensities, apodisation sigma, noise and defocus values are available. Options are also available to toggle Fourier, histogram and noise matching as well as convenient switching to an equivalent fluorescence kernel.

The interactive notebook pictured is available as an example, along with sample images at https://github.com/georgeoshardo/SyMBac/blob/main/examples/Drawing_Phase_Contrast_100x_oil.ipynb. As per the documentation, the adjustable parameters are given by:

- `media_multiplier` is the intensity multiplier for the media part of the image.
- `cell_multiplier` is the intensity multiplier for cell parts of the image.
- `device_multiplier` is the intensity multiplier for the device part of the image.
- `sigma` is the radius (in pixels) of the gaussian apodisation of the phase contrast PSF (if you are using phase contrast).
- `scene_no` is the index for the frame of the synthetic images you rendered.
- `match_fourier` controls whether you are matching the rotational Fourier spectrum of the synthetic image to the real image.
- `match_histogram` controls whether you are matching the intensity histogram of the images with each other.
- `match_noise` controls whether you are matching the camera noise of the images with each other.
- `noise_var` controls the variance of the shot noise added to the image.
- `fluorescence` controls whether you are rendering a fluorescence of phase contrast image.
- `defocus` controls the radius of a gaussian which simulates depth of focus and out of focus effects of the PSF.

### 6 Model Training

Models can be trained with DeLTA [**?**] or Star/Splinedist [7, 8, 9], both of which operate with a U-net at their core. For very low resolution images, Star/Splinedist perform well due to their ability to separate masks from a semantic segmentation to an instance segmentation. This is especially important at low resolutions, where pixel sizes are large and masks may be highly connected. Thus cutting masks for two cells would induce large error in the size measurement. Star/Splinedist’s instance segmentation allows for touching masks with different labels.

However at high magnification (>40x), we found that DeLTA performs far better, and produces extremely accurate masks. One of DeLTA’s features is that it implements the weightmap described in the original U-net paper [10] which is often neglected. This causes the network to pay particular attention to edges where cells are near to one another and touching. We provide interactive notebooks to preprocess, train, and segment data with DeLTA, and we thank the Dunlop Lab, Boston University, for permission to include DeLTA in SyMBac’s codebase.

### 7 Model Evaluation and Segmentation Precision

Figure 2b shows the error’s decrease as training progresses, but the model can overfit. Therefore automated analysis of the segmentation errors (details in subsequent figures) is used to find the epoch which minimises the error. The growth profile of the mother cell lends itself well to automated detection of segmentation errors. The log transform of the length vs time traces out a saw-tooth wave with variable amplitude and phase. By taking the numerical derivative of this data one can find over and under-segmentation errors by simply searching for peaks over a certain threshold and with a certain prominence. These peaks can then be mapped back to the original length trace and the errors corrected with a variety of signal processing techniques.

**Figure SI 11:**
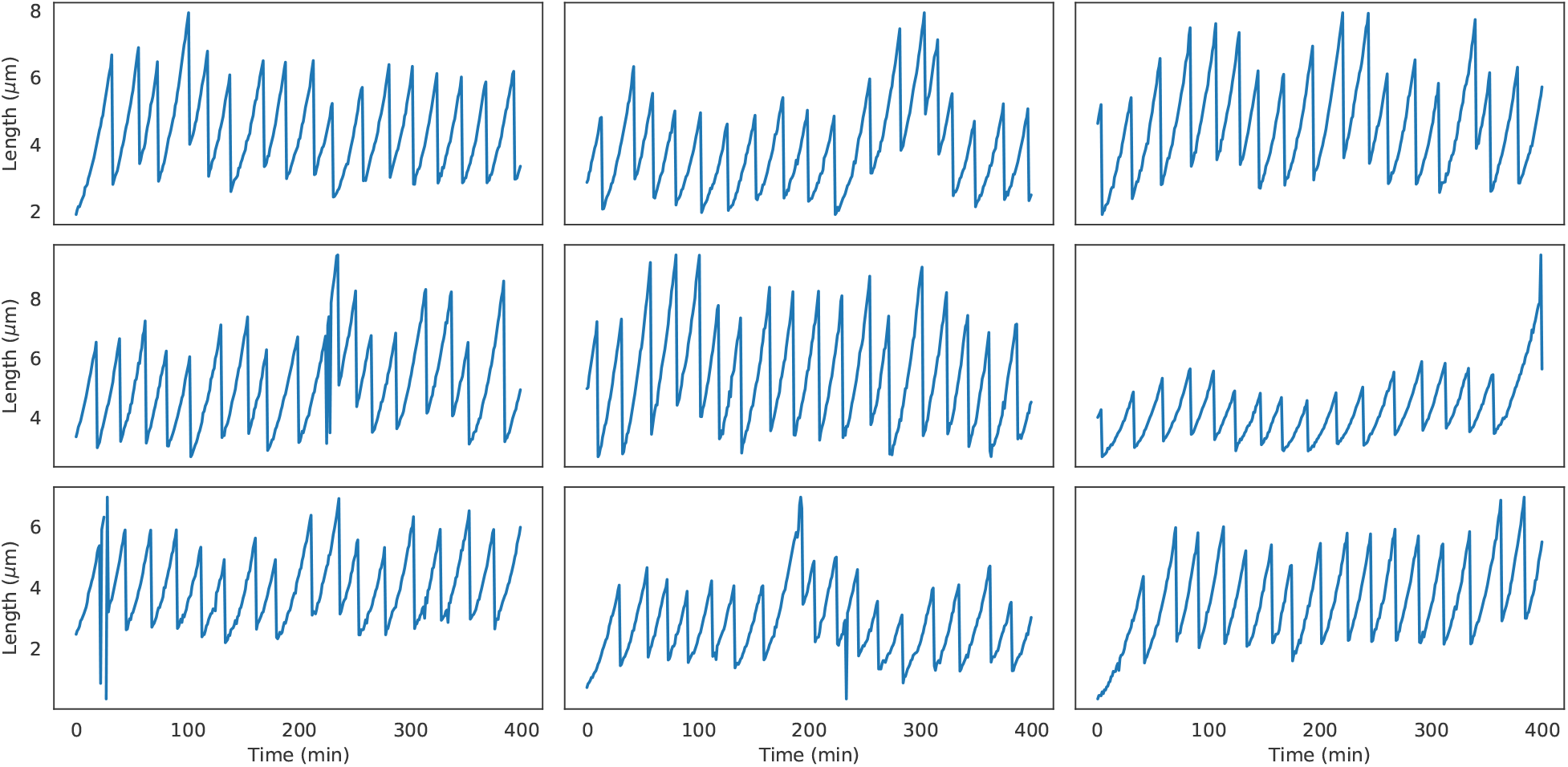
A sample of mother cell growth profiles, reminiscent of saw-tooth waves. (Segmented from 100x oil data, kymographs in Figure SI 16 and Figure SI 17

**Figure SI 12:**
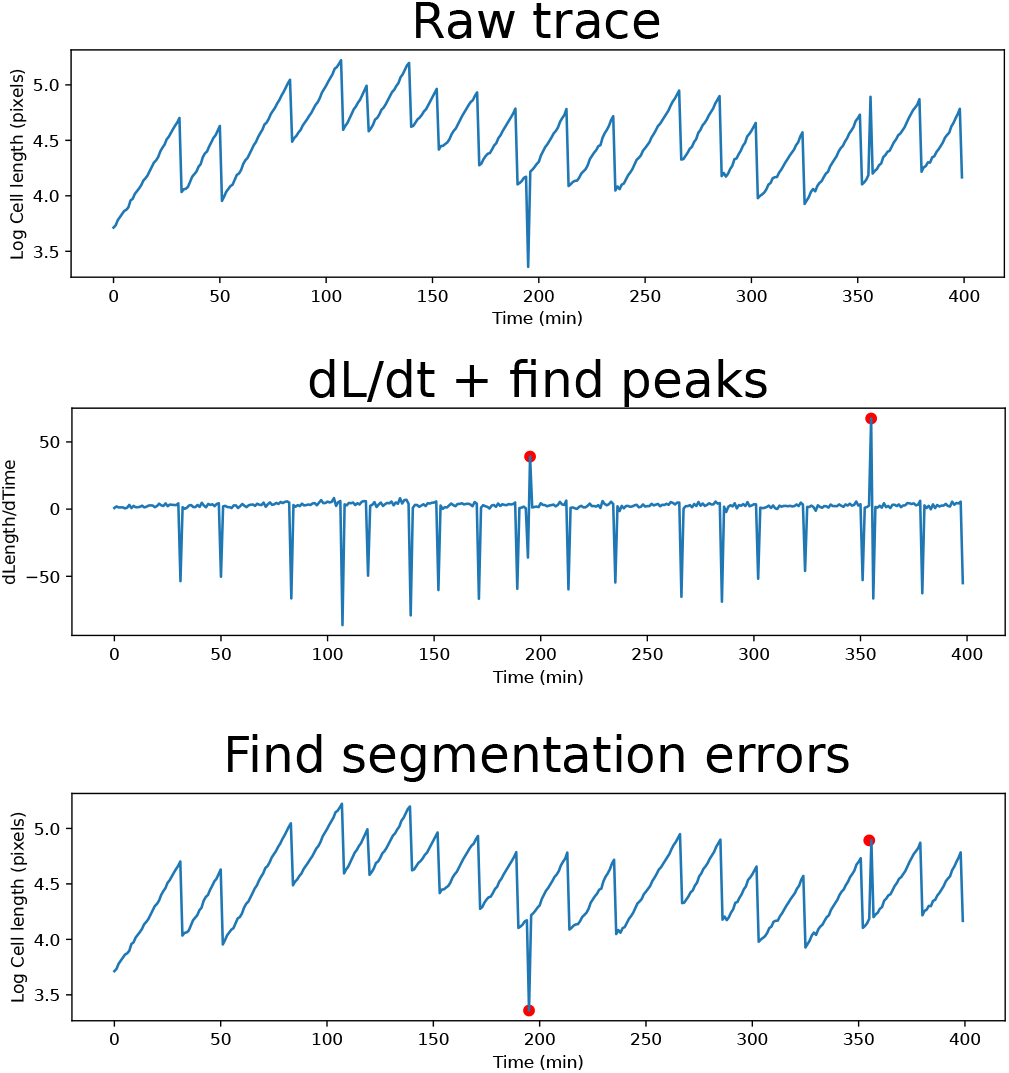
Errors in the segmentation output’s raw length trace are identified by taking its numerical derivative. Peaks in the numerical derivative are found and mapped back to the original data. In this case the errors are fixed simply by smoothing out peaks by setting them to the midpoint value of the adjacent values. For errors which span more than a single frame, errors can still be identified in this way, but require more sophisticated correction methods (for example rolling back by more than one frame, rolling forward by more than one frame, and interpolating between the frames).

Full traces reveal good temporal coherence, and the ability to accurately resolve width over time.

**Figure SI 13:**
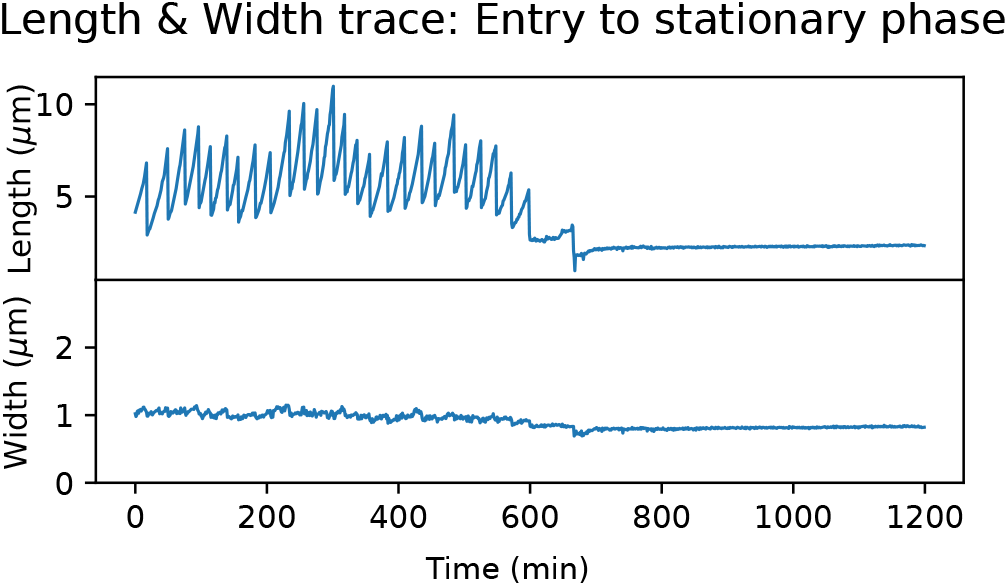
An example trace of a mother cell entering stationary phase. As can be seen from the width and length plots during the stationary phase, there is very high temporal coherence, and the width can be studied at the sub-micron level. At stationary phase, the variance in the width was 100nm^2^, and thus a precision of 10nm can be reached in the estimation of the cell width.

We show that we can achieve a precision of as low as 6.8 nm in the estimation of a cell’s width, and a precision of 20 nm in the estimation of a cell’s length in stationary phase (Figure SI 14). This is only possible due to the high quality training data fed to the model. This is proven by the comparison between human-made and synthetic training in Figure 2g.

**Figure SI 14:**
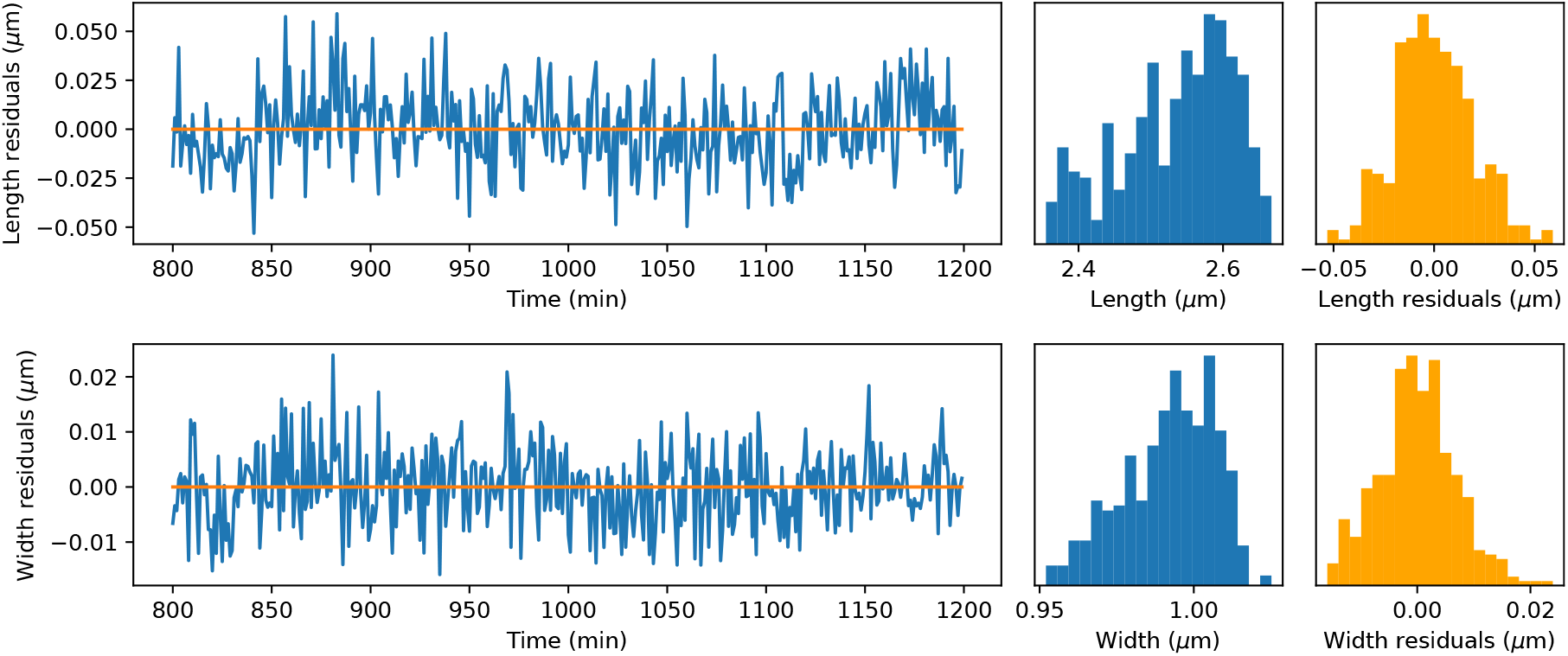
In order to estimate the precision of width and length estimations in the stationary phase, we fit a quadratic polynomial to cell lengths and widths in stationary phase over time. As an example, we show the histograms of the true lengths and widths of this cell, along with histograms of the fit residuals. The standard deviation in the width residuals was 6.8 nm and the standard deviation in the length residuals was 20 nm.

### 8 Temporal Coherence of SyMBac trained models

In order to evaluate the quality of masks generated by models trained on human generated data and computer generated data, we compared the masks outputted by the pretrained model supplied with the DeLTA paper on its own test data, and a model we trained on synthetic data. We first made qualitative observations of mask quality, and then quantified these by assessing the temporal coherence of single cell width between frames. The results of this comparison can be seen in the histogram in Figure 2g, whereby the distribution of output mask widths is tighter for SyMBac trained models. This is further exemplified by noting that the temporal coherence of mask widths was poor as shown below. An example trace of cell widths is shown as a comparison of the outputs from the two types of training data in Figure 2g.

**Figure SI 15:**
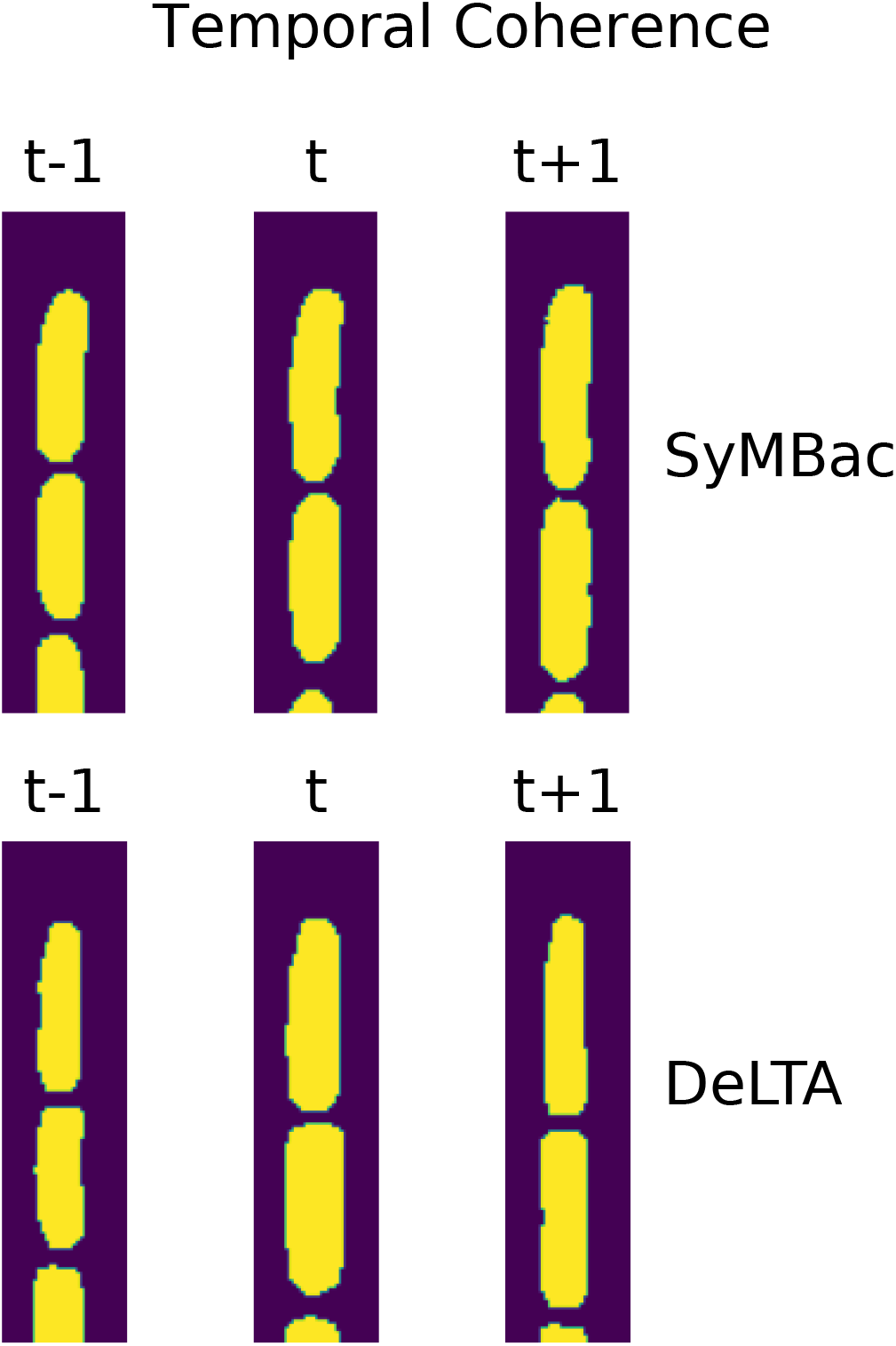
Masks generated from models with SyMBac data can be seen to produce tighter mask width distributions, the result of which is reduced artefactual fluctuations in mask width, leading to higher temporal coherence. The output masks of models trained on the DeLTA training data show large and visible fluctuations in width from frame to frame, as exemplified.

### 9 Segmentation Examples: Kymographs

#### 9.1 Exponential growth (100x oil)

**Figure SI 16:**
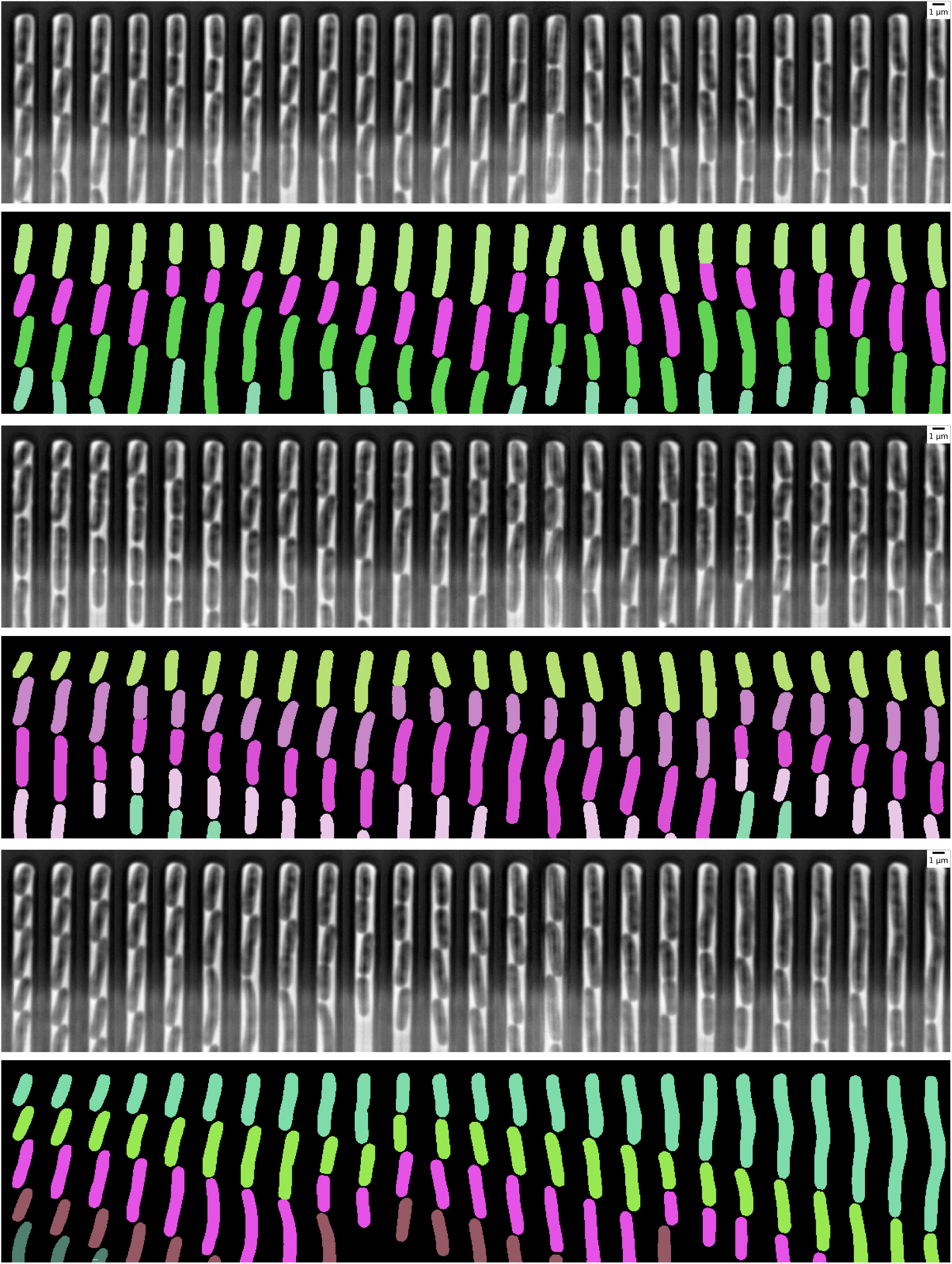
Exponential growth in the mother machine, from Bakshi et al. [11]. Frame spacing is 3 minutes

#### 9.2 Entry to stationary phase (100x oil)

**Figure SI 17:**
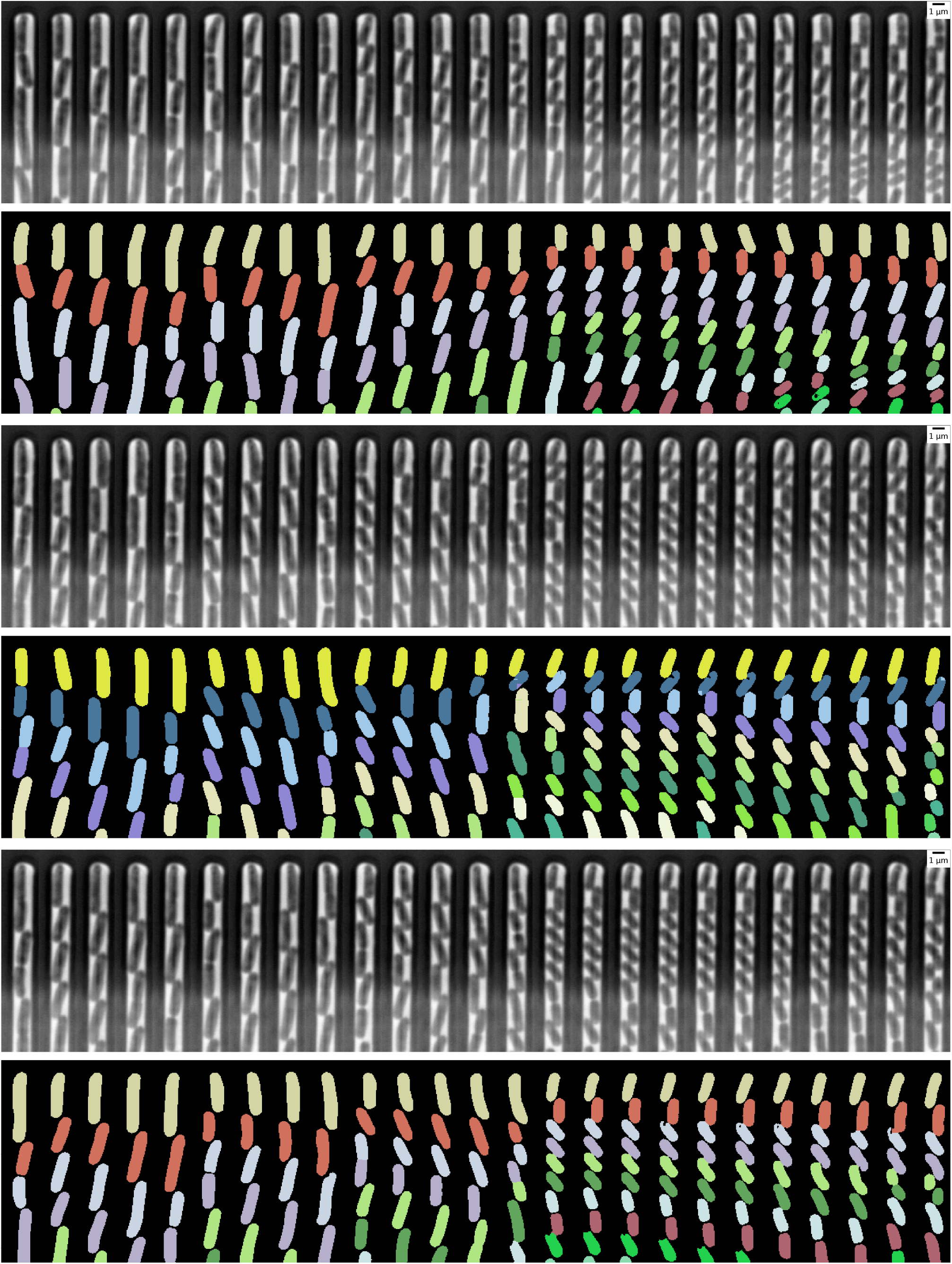
Entry to stationary phase growth, from Bakshi et al. [11]. Frame spacing is 6 minutes

#### 9.3 Exponential Growth (60x air)

**Figure SI 18:**
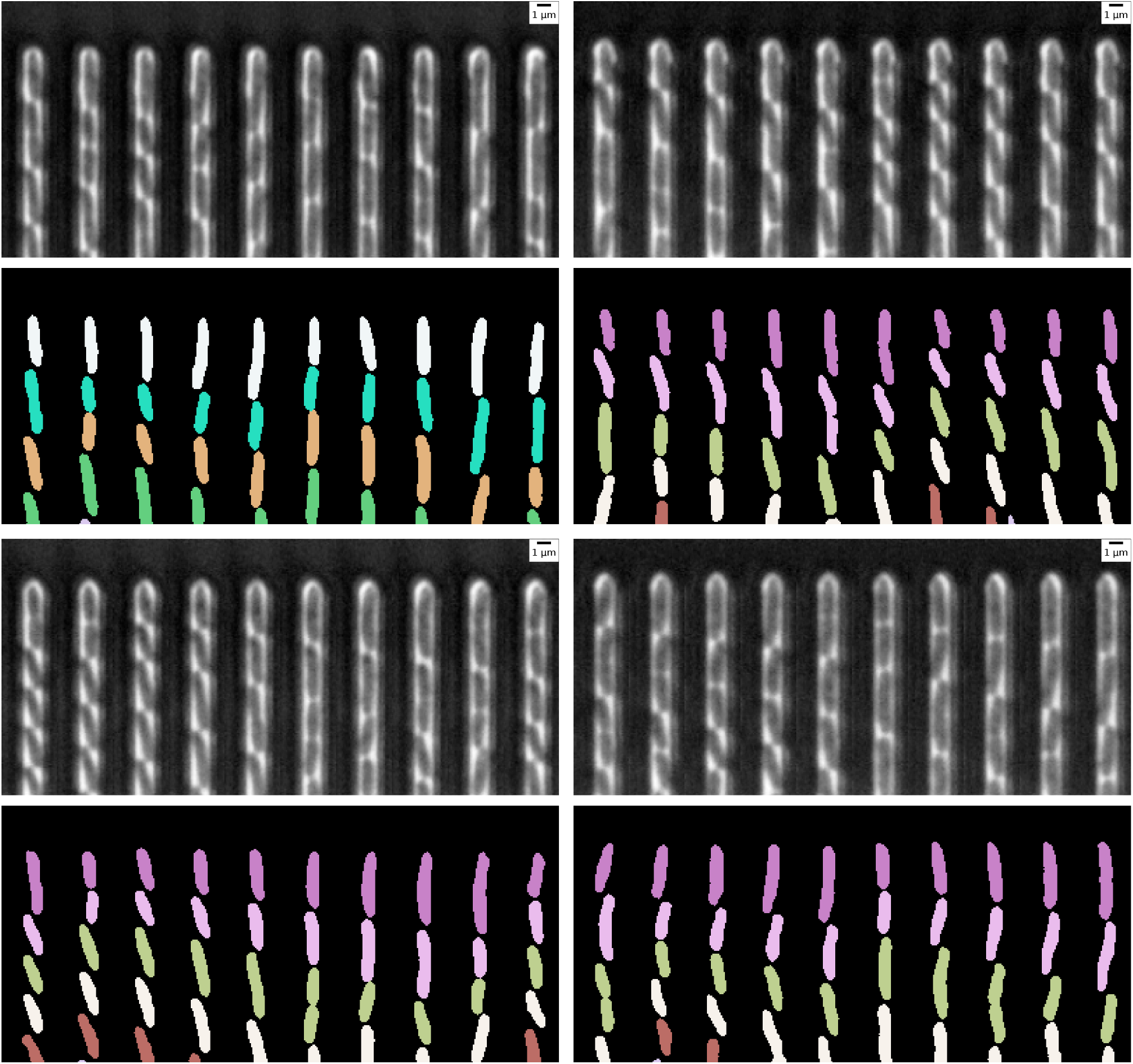
Exponential growth in the mother machine, from Bakshi et al. [11]. Frame spacing is 5 minutes

### 10 Size regulation analysis during exit from stationary phase

Correlations between sizes (area), length, and width at the point of exit from stationary phase and at first division indicates cells are sizers. Irrespective of the initial size, length, width, cells only divide after reaching a critical value.

**Figure SI 19:**
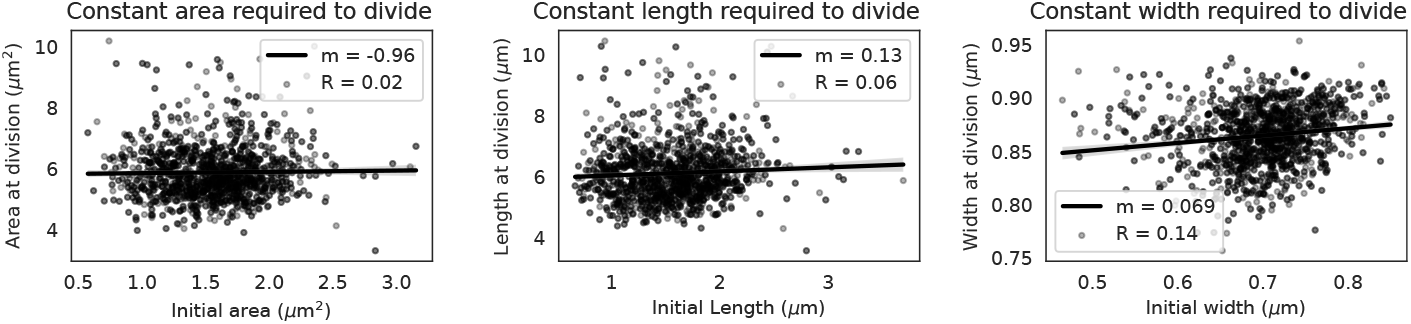
The cells are sizers in every aspect, area, width and length, during exit from stationary phase.

### 11 Extension to 2D growth regimes and other microfluidic geometries

SyMBac works in much the same way for 2D growth as it does for 1D mother machine growth, with a few key differences. Firstly, the cell simulator back-end was switched from our custom simulator (built only with 1D growth in mind) to the more general CellModeller [12] (the reason for not using CellModeller for all simulations, is that CellModeller cannot vary cell width during a simulation, whereas our simulator can, allowing us to capture a variety of widths in the 1D growth data, where it matters most). Simulations of 2D growth (standard agar pad experiments) and channelled 2D growth like that seen using the microfluidic device in [13] were generated, and the cell properties saved. SyMBac was then used to redraw the scenes and apply filters to produce the synthetic data.

#### 11.1 2D microfluidic growth chamber (microfluidic turbedostat)

Microfluidic devices need not be restricted to 1 dimensional growth. Increasing the width of a mother machine trench turns it into a microfluidic chemostat (Figure 1f). Simulating cells in this geometry consists of simply adding two constraints to side of the simulation much in the same way as mother machine growth is simulated. Cells are then removed when they reach the horizontal nutrient flow. Due to the large halo often seen near the flow channel making cells difficult to see, we crop the image to only include the main region of growth. The generated data is then fed into a segmentation network, and an example montage of the output masks is shown in Figure SI 20.

**Figure SI 20:**
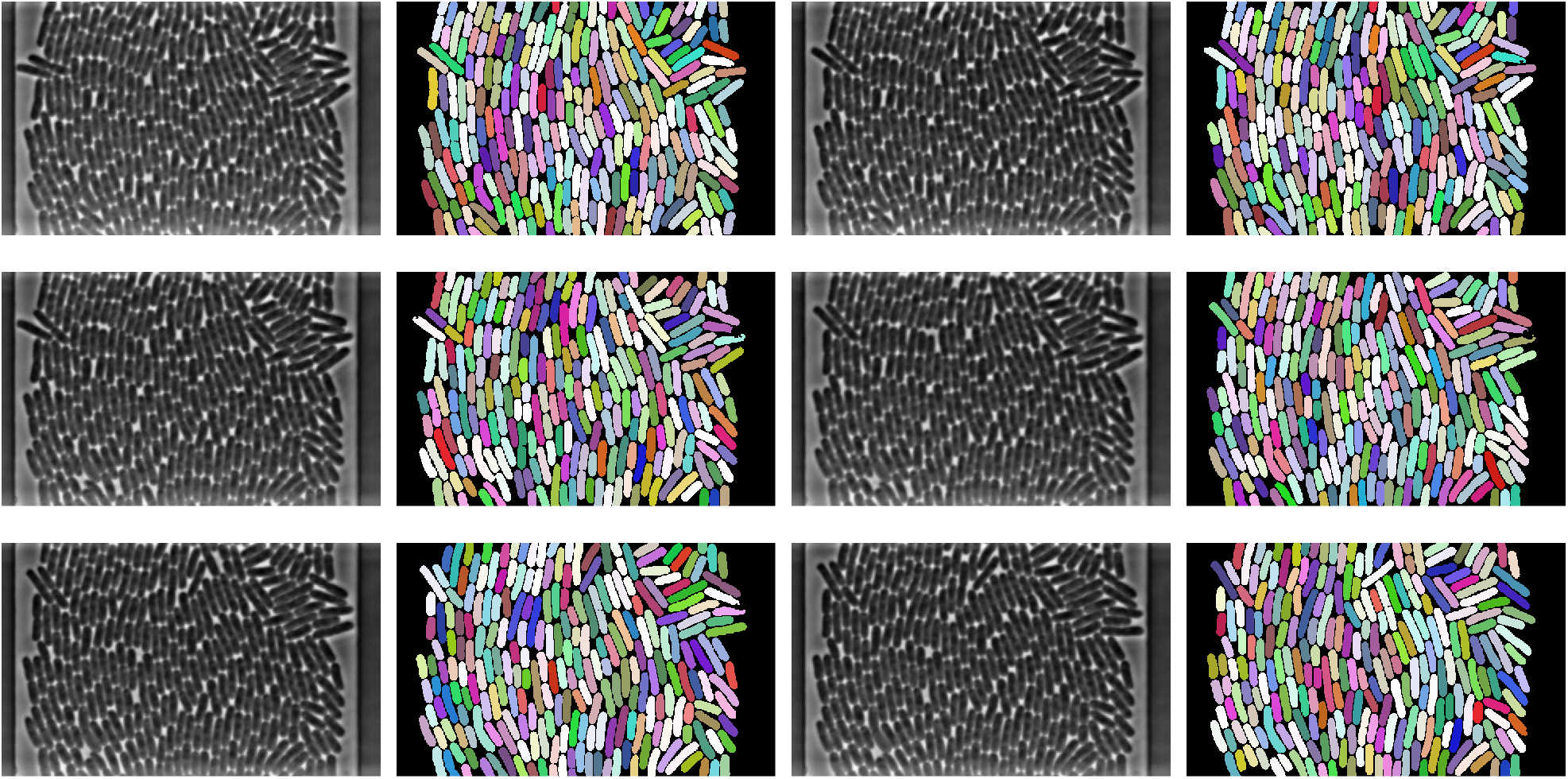
Montage of segmentation outputs from a model trained on SyMBac data, from various sequential timepoints of data supplied by the Elf Lab, Uppsala University.

#### 11.2 Simulating growth of 2D colonies on agar pad

##### 11.2.1 Typical Phase Contrast Image Features in Images of Monolayer Colonies on Agar-pad

Phase contrast images of bacterial growth on agar pads are typically relatively low contrast, with additional texturing/patterning due to the characteristics of the solid agar medium on which the cells grow. Furthermore, due to the interference effects from the phase contrast point spread function, the cells at the colony’s perimeter are darker, surrounded by a bright halo at the interface between the medium and the cell. Another phase contrast artefact is the shade-off, which bleeds into the centre of the sample causing internal cells to be lighter in colour. While these artefacts are not easily seen in images of cells in linear colonies (e.g. mother machine images), they are very visible in almost all agar pad experiments. Our phase contrast image generation pipeline (described in Equation 3) was augmented with the addition of a very small offset to the PSF (dependent on the size of the kernel) which can be used to precisely modulate the amount of halo and shade-off in any given synthetic image. We found that a rough initial guess with random sampling around it was sufficient to generate very high quality training data capable of training highly accurate U-net models.

**Figure SI 21:**
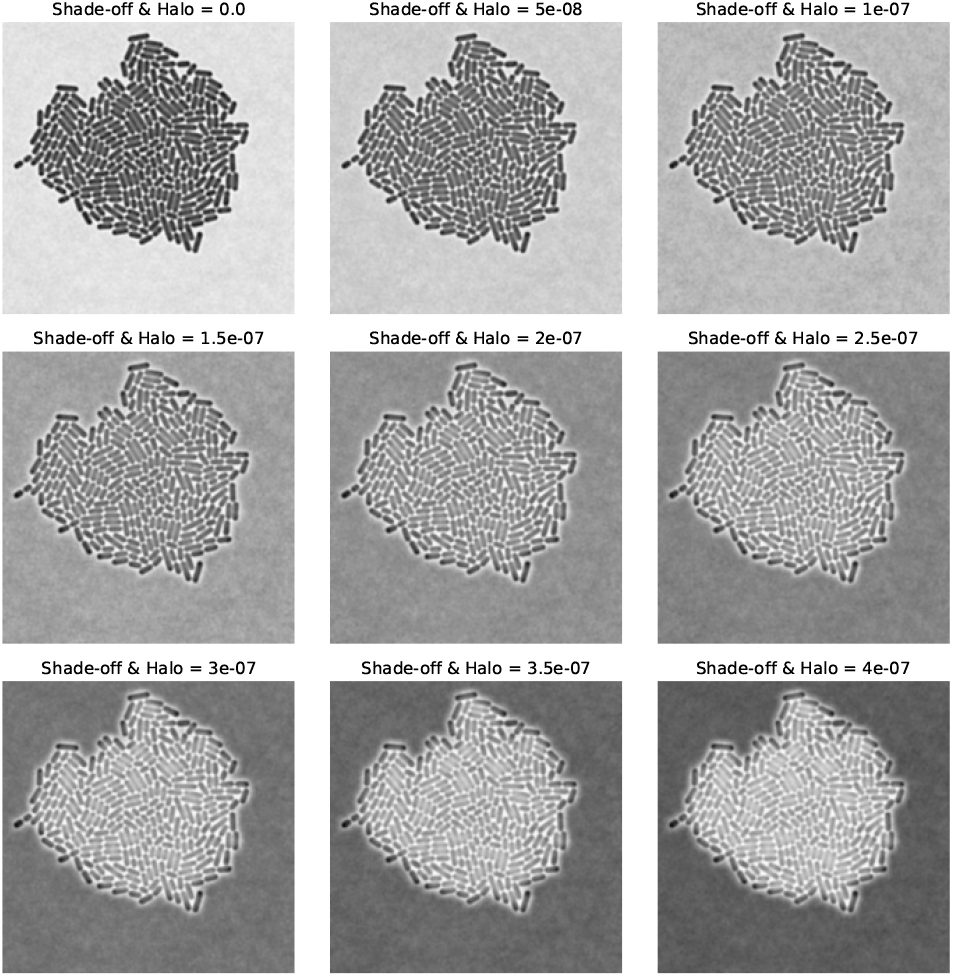
Increasing shade-off and halo phase effects with increasing PSF offset.

Additionally, phase contrast agar pad images typically have some form of texturing in the background due to anisotropy in the agar. Perlin noise [14] is often used to generate natural looking surfaces and terrain heightmaps in computer graphics. For this reason we used Perlin noise with a variety of randomised parameters to simulate the defects seen in phase contrast images of agar pads. Examples of this can be seen in Figure SI 22. The addition of this noise greatly increases training accuracy, as without it background noise is often positively segmented, creating spurious phantom masks.

**Figure SI 22:**
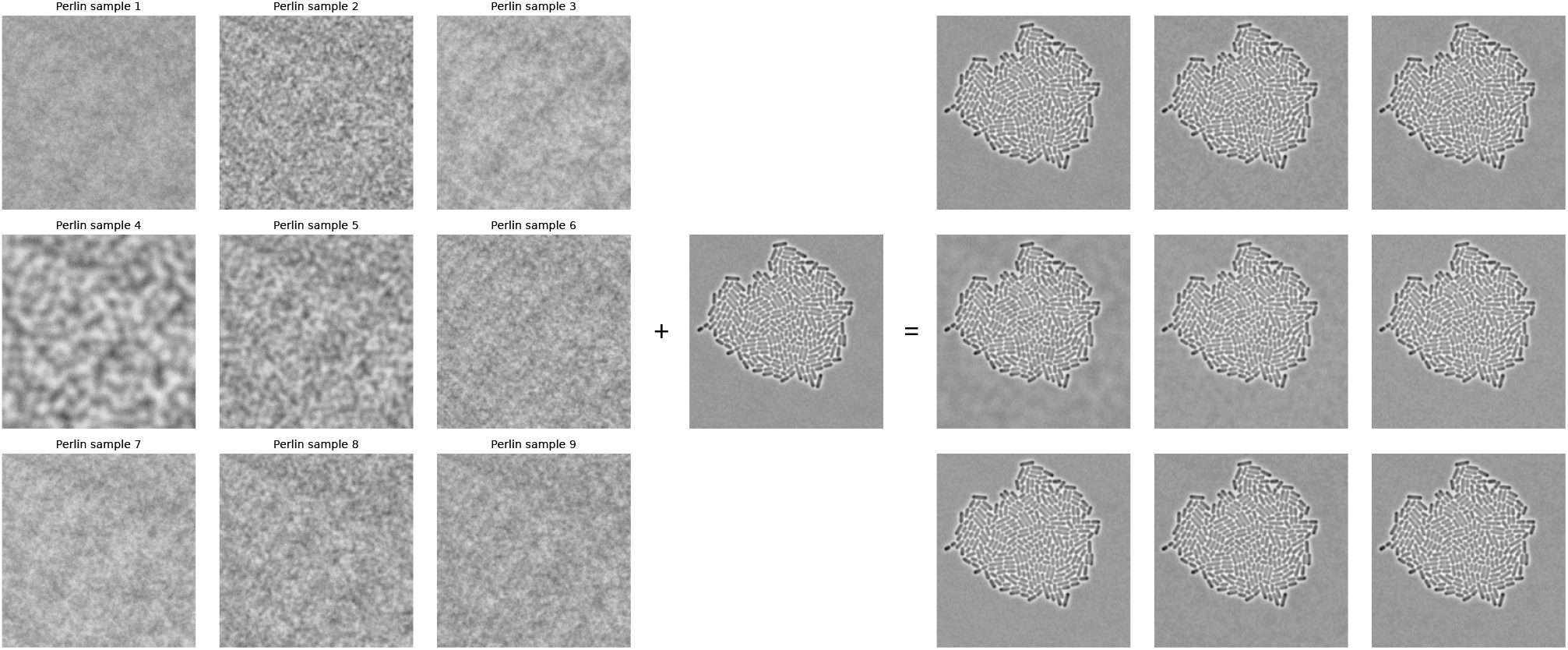
Random sample of Perlin noise (shown on the left with enhanced contrast) made to mimic agar pad phase defects can be incorporated with synthetic images to create more realistic agar backgrounds.

To test the efficacy of this training data, we analysed bacteria growing in a monolayer on agar pads. An example montage of the segmentation is shown below. The performance on this data is very good considering no post-processing is done other than a simple threshold on the U-net probability output.

**Figure SI 23:**
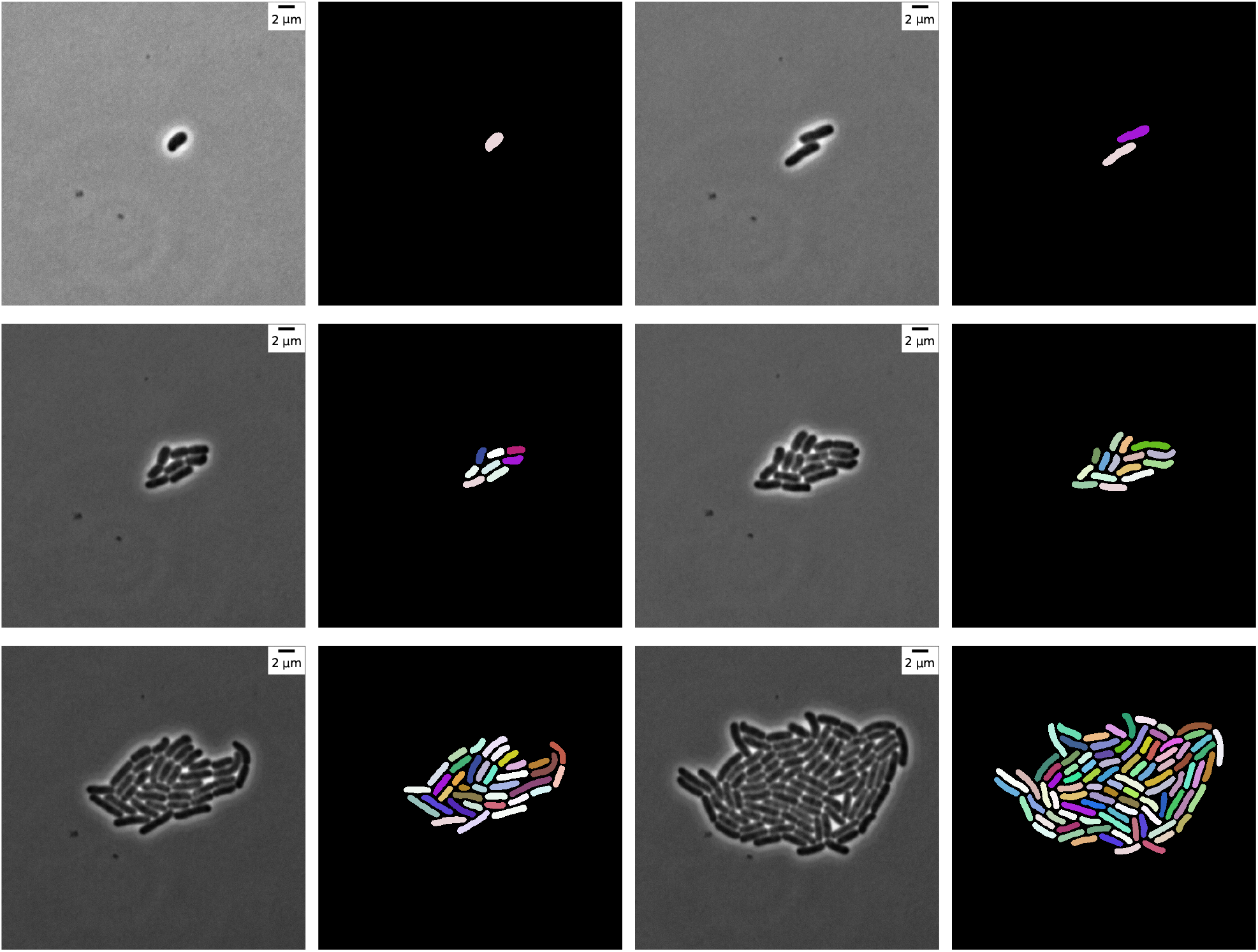
Montage of *E. coli* growing on an agar pad, segmented using a U-net trained on SyMBac synthetic data.

##### 11.2.2 Fluorescence Images of Monolayer Colonies on Agar Pad

Generating fluorescent data is identical to the generation of phase contrast data (described above), except we use a fluorescence PSF, and do not add any pseudorandom noise to the image background. The only noise added to the image is shot noise, which simulates the camera’s noise (used in all image generation schemes). Additionally, a large variability was added in the brightness of each cell to account for real changes in fluorescent intensity due to stochastic gene expression. To test the ability of our synthetic data to train a model, we streaked a dense culture of stationary phase *E. coli* expressing YFP onto an agar pad and imaged a large field of view with a 60x air objective. These images were very large (1200×1200 pixels), and so synthetic training data of size 400×400 was generated (shown in Figure SI 24), and the image was segmented patch by patch. Examples of segmented FOVs are shown below in Figure SI 25.

**Figure SI 24:**
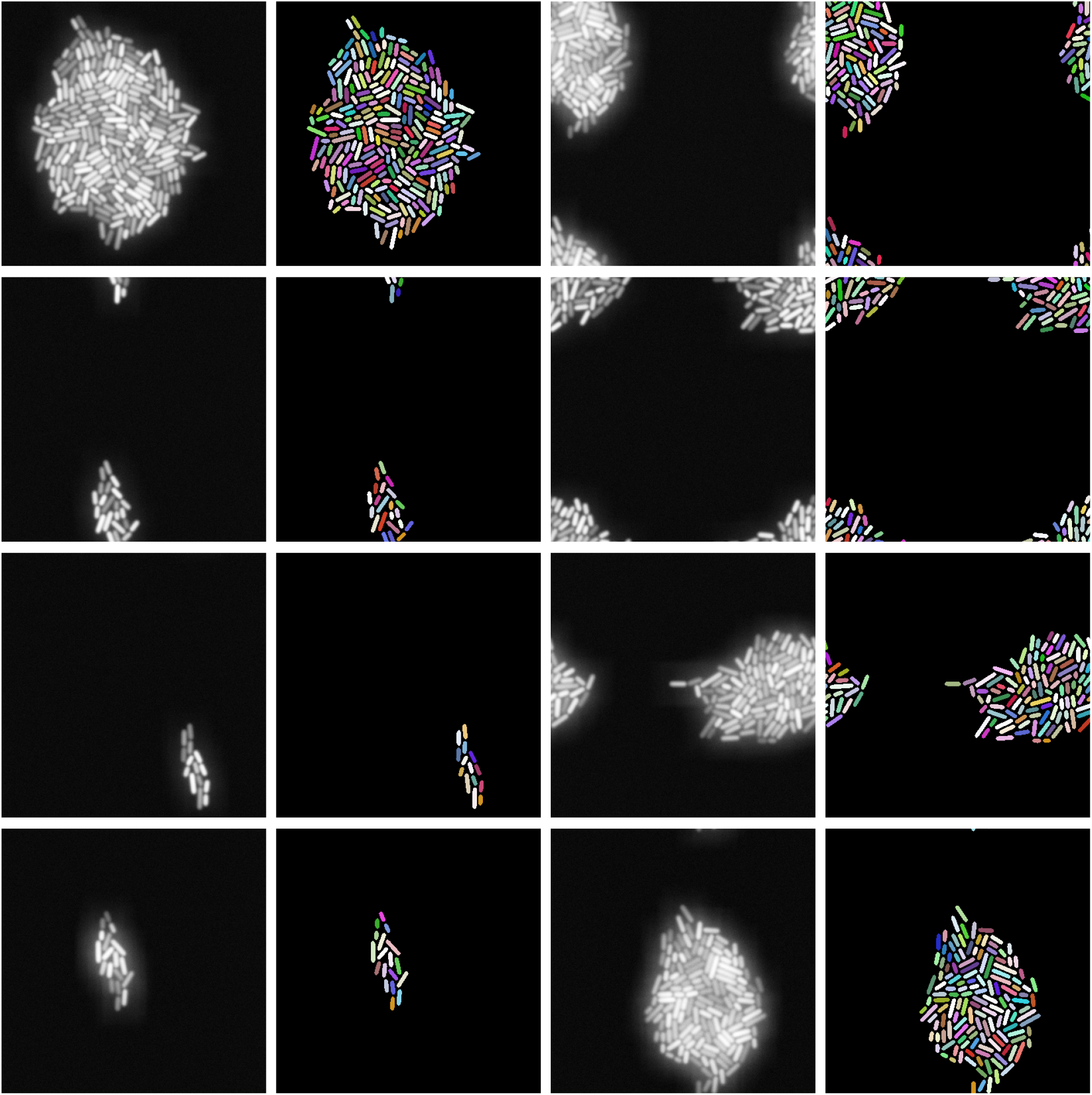
Synthetic fluorescence training samples with accompanying masks.

**Figure SI 25:**
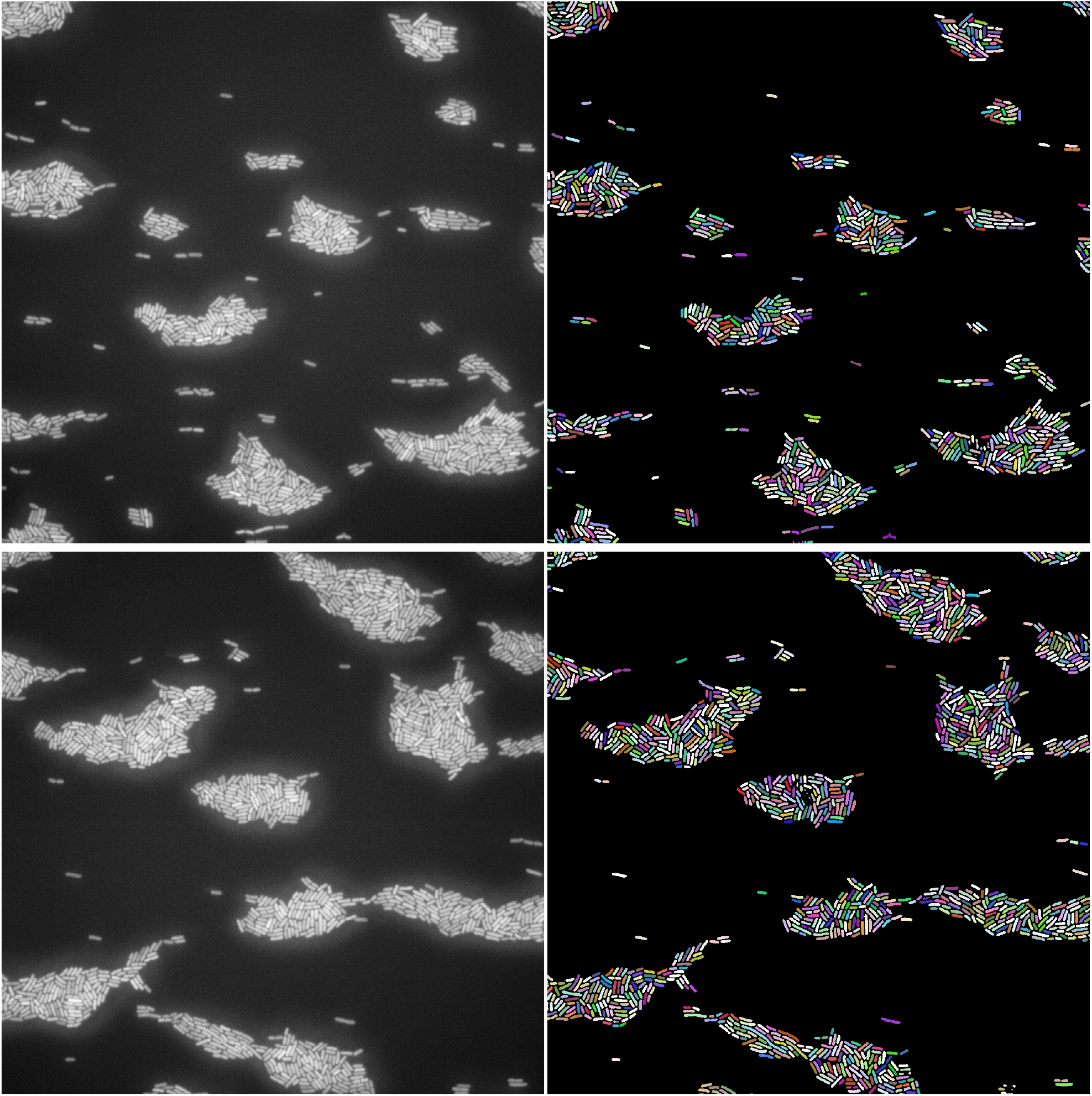
Whole FOVs segmented patch by patch on a model trained on synthetic fluorescence data.

